# Beyond Sequence and Structure: Pretraining PLMs on Continuous Biophysical Fields

**DOI:** 10.64898/2026.11.09.765464

**Authors:** Patrick Jiang, Abhishek Singharoy

## Abstract

The Biophysical Transformer is a protein language model that adds continuous biophysical fields as a third pre-training signal beside sequence and structure. A Poisson–Boltzmann solver computes the electrostatic potential around each three-residue token, and the model learns to predict it from sequence alone. A rotationally invariant spherical-harmonic descriptor serves as the loss target, eliminating the need for frame alignment during training and for coordinates at inference. On FLIP Meltome thermostability, the field objective raises Spearman correlation against a matched, one-flag control, with gains that survive residualizing dipeptide composition and removing overlapping test proteins. On the CAID disorder benchmark, the field arm improves AUC-ROC from 0.73 to 0.76 and MCC from 0.31 to 0.37 over the sequence-only control—with only 5% pretraining overlap versus 91% for ESM-2. On PLM-Interact protein–protein interaction prediction, the field objective raises AUPRC on two of three tasks (+0.087 cross-species, +0.082 virus–human) at 3–4% corpus overlap. On MHC binding prediction, the 600×-smaller BioTransformer outperforms ESM-2 outright (Spearman 0.76 vs. 0.64 on MHC I, 0.61 vs. 0.52 on MHC II). Across four benchmarks, a 1.1M-parameter model pretrained on 21K structures consistently benefits from explicit electrostatic supervision, establishing continuous physical fields as a viable and complementary pretraining signal for protein language models.

## 1. Introduction

Protein language models learn from sequences, and structure predictors learn from coordinates. Both signals are discrete: sequences are strings over a 20-letter alphabet, and structures are lists of atomic positions in an arbitrary crystallographic frame (Lin et al., 2023; Jumper et al., 2021). Yet neither directly represents the continuous physical environment a protein creates. Quantities like the electrostatic potential, the solvation field and the shape of the binding surface are all functions of the coordinates, but no coordinate list states them explicitly.

We introduce the *Biophysical Transformer*, a protein language model that receives a third pretraining signal: the electrostatic potential computed around each residue by a Poisson–Boltzmann solver (Figure 1a). During training, the model learns to predict this field from sequence alone, as an auxiliary objective alongside masked language modeling. Because the sequence does not determine the orientation of a deposited structure, we score a rotationally invariant descriptor, built from spherical-harmonic shell statistics, so that training requires no frame alignment and inference requires no coordinates (Figure 1b).

**Figure 1.**
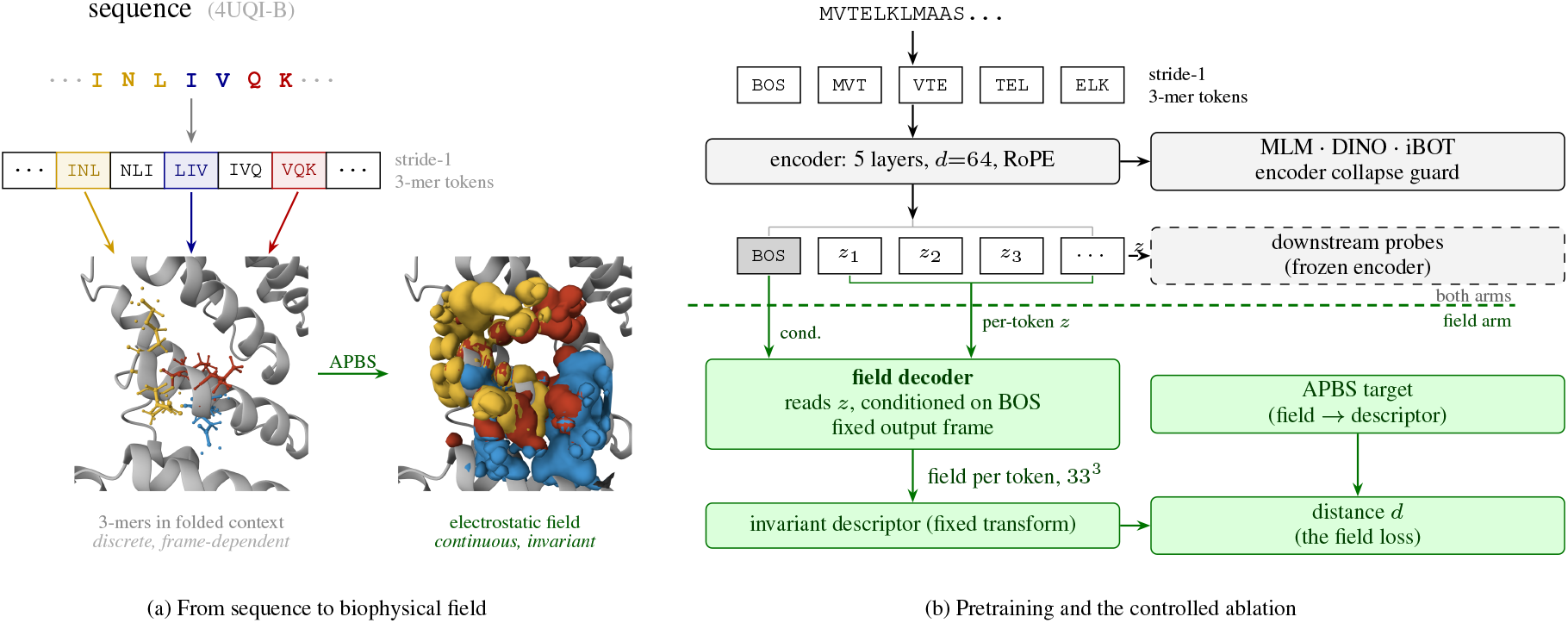
A third supervision signal, and the experiment that measures its effect. *(a)* A selected 3-mer in the amino acid sequence (*top*), its backbone in the folded protein (*left*), and the environmentally aware electrostatic potential computed by APBS around the same residues (*right*). The sequence is discrete and one-dimensional; the coordinates are discrete and frame-dependent; the field is continuous and, through a spherical-harmonic descriptor, invariant to the crystallographic frame. *(b)* Both arms train a 3-mer encoder with MLM, DINO and iBOT. In the field arm a decoder reads the per-token states *z*, conditioned on the BOS state, emits a field per token in a fixed frame (Figure 5), and is scored against the APBS target through the invariant descriptor. One flag removes everything below the dashed line, leaving a sequence-only control identical at initialization. Green marks the field arm alone; probes read *z* in both arms.

**Figure 2.**
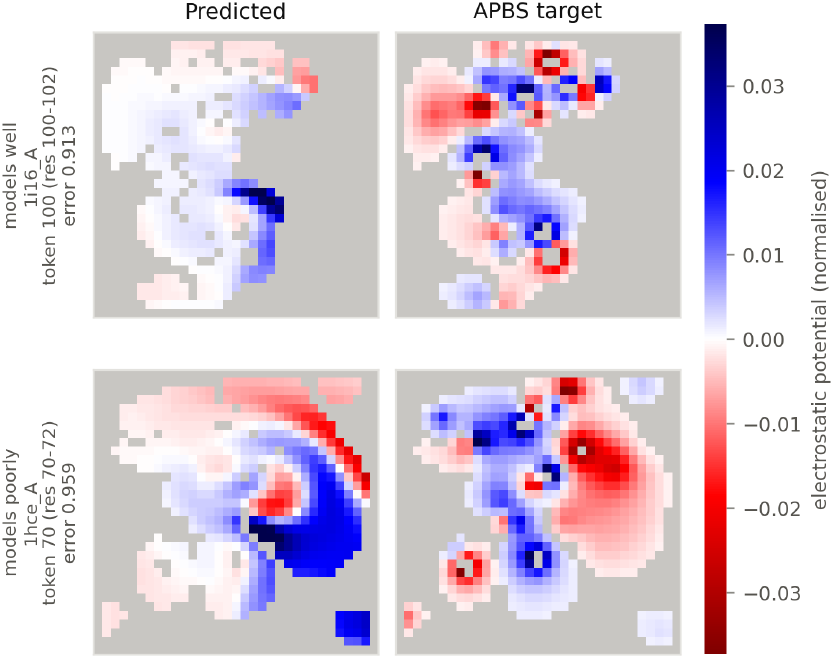
Decoder output beside the APBS target for two heldout proteins. Mid-plane slices of the 33^3^ box. Rows are the 10th and 90th percentile proteins of masked-token perplexity among 293 held-out proteins (1i16_A token 100 and 1hce_A token 70). Each shows the token whose descriptor error is closest to its protein’s median. Quantitative reconstruction accuracy is in Appendix J.

**Figure 3.**
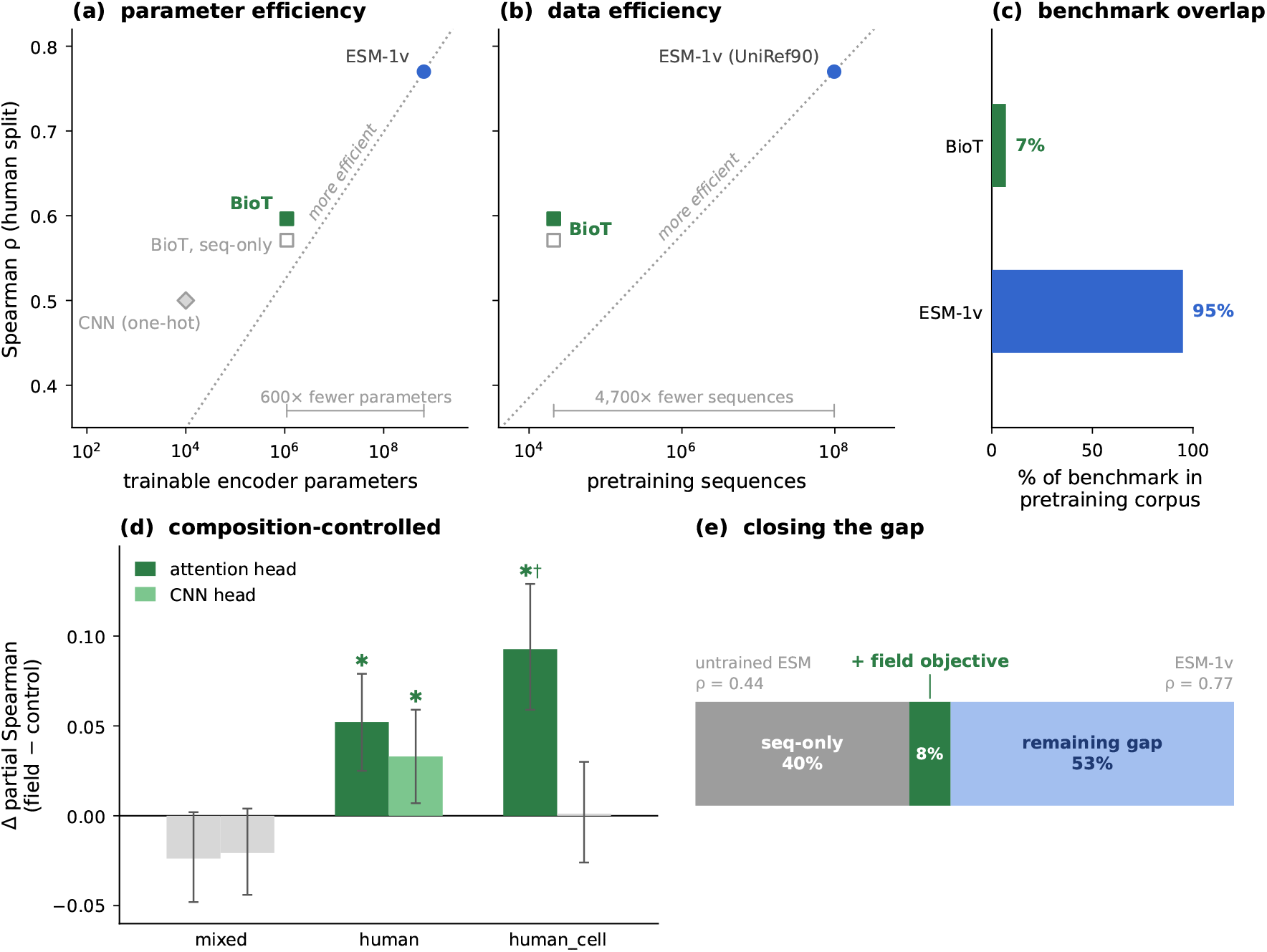
Resource budget, benchmark exposure, and the controlled effect of the field objective. *(a)* Spearman *ρ* on the human split versus trainable encoder parameters; the Biophysical Transformer’s 1.1M-parameter field arm sits 600 *×* below ESM-1v yet closes nearly half the gap from an untrained 650M encoder. *(b)* The same comparison on the data axis: 21K structures versus 98M sequences. *(c)* Fraction of FLIP Meltome human test proteins inside each model’s pretraining corpus; ESM-1v covers ∼95% through UniProt membership, the Biophysical Transformer’s 7% at the 0.80 3-mer threshold (Section 5.3). *(d)* Δ partial Spearman (field minus control) after residualizing 400-dimensional dipeptide features. Stars mark 95% intervals excluding zero; *†*marks an estimate unstable under protein removal (Section 6). *(e)* Decomposition of the untrained-to-ESM-1v gap on human: the sequence-only arm recovers 40%, the field objective adds 8%, and 53% remains.

As a deterministic function of the coordinates, the field carries no information a structure lacks. What it contributes instead is the physical model’s own assumptions (continuum electrostatics, a fixed charge model, solvation parameters) as a structured inductive bias that a small model might not discover from sequence co-occurrence alone. Evaluated on the FLIP Meltome thermostability benchmark (Dallago et al., 2021) with frozen embeddings, the field objective raises Spearman correlation on both human splits against a matched, one-flag control. The gain on `human` survives residualizing dipeptide composition under two architecturally different readouts and survives removing the test proteins that overlap the pretraining corpus. No gain survives on the organism-mixed split, however.

Our contributions are (i) the invariant field objective (Section 3.3), (ii) a controlled ablation in which both arms are identical at initialization and diverge only through the field loss (Section 4), and (iii) transfer experiments on intrinsic disorder, protein–protein interaction and MHC binding that test whether the field signal generalises beyond the primary benchmark. All results rest on a single pretraining run per arm, and the design cannot yet separate field content from generic regularization (Section 6).

## 2. Related Work

Large-scale masked language modeling on protein sequences has shown that co-evolutionary statistics encode substantial structural and functional information. ESM-2 demonstrated that a single sequence model, trained on hundreds of millions of UniRef sequences, can predict atomic-level structure from its internal representations alone (Lin et al., 2023). ProtTrans confirmed these findings across multiple transformer architectures and training objectives (Elnaggar et al., 2022). The MSA Transformer (Rao et al., 2021) showed that co-evolutionary signal from multiple sequence alignments can be distilled into a single-sequence model, and ProstT5 further closed the gap between sequence and structure models by translating between sequence and structure tokens (Heinzinger et al., 2024). These models extract physics implicitly through the statistical regularities of evolution, but none receives an explicitly physical supervision signal during training.

Structure predictors take the complementary route, learning directly from three-dimensional coordinates. AlphaFold (Jumper et al., 2021) and its successors (Abramson et al., 2024; Passaro et al., 2025) achieve near-experimental accuracy on many targets, but their representations end in discrete atomic positions, situated in an arbitrary crystallo-graphic frame. Aligning or canonicalizing that frame is a recurring design constraint. Frame-invariant structural representations have been explored through internal coordi-nates, distance matrices and equivariant architectures, but these still describe geometry with points rather than with the continuous fields that geometry generates.

Volumetric representations of the molecular environment have a history as *inputs* to prediction models. Torng and Altman used 3D CNNs on voxelized amino acid environments for functional site detection and similarity analysis (Torng & Altman, 2017; 2019), and DeepSurf extended this approach to molecular surfaces for binding-site prediction (Mylonas et al., 2021). In each case, the method computes the three-dimensional field or surface from the structure and hands it to the model as input. Our work reverses the direction: the electrostatic potential, computed by the APBS Poisson–Boltzmann solver (Jurrus et al., 2018), serves as a prediction *target* for a sequence-only model.

Concurrent with our work, METL (Gelman et al., 2025) pretrains transformers on Rosetta energy landscapes and fine-tunes them on experimental fitness data, demonstrating that biophysical simulation can replace evolutionary statistics as a pretraining signal for protein engineering tasks. METL’s targets are scalar energy terms per mutant; ours are spatially resolved electrostatic fields per token. The two approaches are therefore complementary: METL captures the global energy surface of a single protein variant, whereas the Biophysical Transformer captures the local electrostatic environment of each residue and aggre-gates it through the encoder. Whether the two signals are additive when combined into one model is an open question.

The descriptor we use to compare predicted and target fields draws on the SOAP formalism for representing chemical environments (Bartók et al., 2013), which decomposes local atomic environments into rotationally invariant spherical-harmonic power spectra. We adapt this idea to voxelized electrostatic fields, decomposing each residue’s field into radial shell coefficients and cross-shell power matrices that are invariant to the deposited orientation.

## 3. Method

### 3.1. Tokenization

Sequences become stride-1 3-mers (*L* − 2 tokens after a BOS token), each carrying the field of its three-residue window. Terminal residues get no field of their own. We chose *k*=3 so that a residue’s covalent neighbors, which dominate the near-field potential, sit inside the token and attention supplies only through-space contributions. It is also the largest *k* the model can afford: a *k*=4 vocabulary would be about nine times the model’s size. We did not vary *k*.

### 3.2. Architecture

The Biophysical Transformer is a PLM with one extra objective. Its encoder is a 5-layer transformer (*d*=64, 4 heads, RoPE positional encoding (Su et al., 2024)), and its pertoken output *z*, BOS included, is what every downstream probe reads. The encoder trains in both arms of the ablation and is the only component any evaluation touches.

In the field arm, a decoder reads *z* and emits a 33^3^ electrostatic potential grid per token in a fixed output frame. The invariant descriptor scores this output against the APBS target (Section 3.3). The key design choice is that non-equivariant projections of the BOS state build a seed volume on a fixed 4^3^ coordinate grid, which anchors the output frame, and SE(3)-equivariant layers then refine it to the full 33^3^ resolution. The loss is indifferent to the output frame, exactly under the 24 cube rotations and approximately under continuous ones (Appendix D). Appendix C gives the full decoder architecture. Of 1.1M trainable parameters, 679K sit in the encoder and 398K in the decoder path.

### 3.3. Invariant Objective

The target is a rotationally invariant descriptor of each token’s field (Appendix D): over 12 radial shells, the *l*=0 shell means (linear in the field, so carrying its sign) and, for each *l*=1, …, 6, the cross-shell power matrix (quadratic, so sign-blind). The angular block is compressed,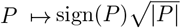, so that its gradient stays order one as the output approaches the spherically symmetric stationary point. The loss scores the seven degree blocks separately and averages them, so only one of the seven blocks supervises polarity (Section 6).

We supervise the descriptor rather than the field because the sequence does not determine target orientation. Let *V* be the field on a grid aligned to the structure’s own frame, and *G* the 24 grid-preserving rotations. If orientation is uniform over *G* and independent of the sequence *S*,

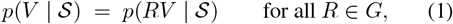

then the risk-minimizing prediction is *G*-invariant (for any strictly convex voxel-wise loss), so it has no dipole, quadrupole or octupole component and loses exactly the angular information that distinguishes residues. A voxelwise variant we trained collapsed in this way: the decoder produced smooth, isotropic predictions indistinguishable across residue types. Uniformity holds only approximately: the corpus-mean field carries a coherent dipole at about 3% of a typical protein’s own (Appendix F). However, this is a single direction shared across the corpus and identical for every *S*, so it distinguishes no residue from another and the argument is unaffected.

Descriptors are compared with

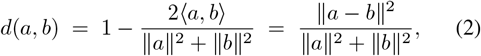

which lies in [0, 2], equals 1 if either descriptor is zero, and is applied per degree block then averaged. Four invariant terms and a masked variant make up the objective (see Appendix E for the terms, Appendix A for weights). The primary term is residue-centered descriptor agreement. The remaining terms pin the protein-wide level, the radial energy and the excess total variation. Masked field reconstruction, the single largest weight in the objective (effective 3.0), scores the field at positions whose residue is absent from the input, so the loss falls only insofar as *z* carries that residue’s environment from context. This is how the field objective reaches the sequence representation.

### 3.4. Field Corpus

Each token carries a 33^3^ potential grid computed from the deposited structure, axis-aligned to its frame. The corpus holds 21,418 training and 5,355 held-out proteins drawn from CATH, clustered first with MMseqs2 (Steinegger & Söding, 2017) then with Foldseek for structural coverage. Potentials come from APBS 3.4.1 (Jurrus et al., 2018) on pdb2pqr 3.7.1 (Dolinsky et al., 2004) PQR files with PARSE parameters (Sitkoff et al., 1994). Solver settings beyond the software versions and parameter set are not reported (Section 6). The held-out set also selects the pre-training checkpoint (Appendix H).

Two features of the corpus shape interpretation. First, it leans human: *Homo sapiens* accounts for 18.1% of proteins against 3.1% for *Mus musculus*, a candidate explanation for the human-specific downstream effect. Second, no identity filter separates training from held-out proteins: 241 (4.5%) of the 5,355 held-out proteins share a 3-mer token set with a training protein. Overlap with the FLIP benchmark is audited in Appendix K.

### 3.5. Training

The field objective trains jointly with MLM, DINO (Caron et al., 2021) and iBOT (Zhou et al., 2022) at fixed weights (Appendices A and B): 80 epochs on 4 × H100 in bf16, effective batch size 16, AdamW (Loshchilov & Hutter, 2019) (*β*_1,2_=0.9, 0.95, weight decay 10^*−*4^), encoder learn-ing rate 10^*−*4^ with the decoder rate set ten times higher, 2000 warmup steps and cosine decay to 0.05 of the base rate. The field objective enters at epoch 1.

## 4. Experimental Setup

The control uses the same code, random seed, data order and batches, with the field objective switched off by one flag; the encoder collapse guard runs in both arms. The arms are identical at initialization (all 352 tensors, 2.93M values, bit-identical) and diverge only through the field loss and its latent collapse guard; every evaluation reads only the encoder output *z* (Appendix G).

We evaluate on FLIP Meltome (Dallago et al., 2021) (Spearman correlation) over the mixed, `human` and `human_cell` splits, with 24,817, 8,148 and 5,792 train-ing and 3,134, 1,945 and 1,366 test proteins respectively. FLIP derives the three splits from the Meltome Atlas (Jarzab et al., 2020) by clustering sequences with MMseqs2 (Steinegger & Söding, 2017) at 20% identity and testing on cluster representatives: mixed spans all 13 species, `human` uses human proteins, and `human_cell` one human cell line. FLIP’s released mixed training set differs from the whole-cluster set its text describes (FLIP contributors, 2022); we use the released files. We compute embeddings once from *z* and cache them. Only the readout heads train.

Readouts comprise two composition baselines (a 23-feature amino acid vector and the 400 dipeptide fractions, identical in both arms and therefore a check on the harness), three ridge probes (mean-pooled, BOS, summary statistics), and two trained heads (multi-kernel convolution and gated attention pooling). Trained-head numbers are means over 10 paired seeds; we evaluate each arm at its own best-validation checkpoint (Appendix H).

## 5. Results

### 5.1. The Field Objective Improves Human-Split Thermostability

Adding the electrostatic field objective to a sequence PLM improves downstream thermostability prediction on human protein splits (Table 1). On `human`, both architecturally different trained heads show gains that are two to three times their seed noise: +0.0243 for the CNN head and +0.0253 for attention, each with 95% bootstrap intervals clear of zero and seed standard deviations below 0.007. The agreement of two independent readout architectures reading the same frozen encoder makes an artifact of either head unlikely.

**Table 1.** FLIP Meltome, Spearman. Δ is field minus sequence-only, computed before rounding, with a 95% paired bootstrap interval over test proteins (no seed or pretraining-run variance; one run per arm). Bold marks intervals excluding zero. Trained-head seed s.d. is at most 0.016; full results and composition baselines are in Appendix L.

| Split | Probe | Seq.-only | + Field | Δ [95% interval] |
|---|---|---|---|---|
| Mixed | Ridge (mean) | 0.3496 | 0.3718 | **+0.0222 [+0.001, +0.044]** |
|  | Ridge (BOS) | 0.2810 | 0.2966 | +0.0156 [−0.009, +0.040] |
|  | Ridge (summ.) | 0.3717 | 0.3876 | +0.0159 [−0.006, +0.039] |
|  | CNN head | 0.4371 | 0.4369 | −0.0002 [−0.019, +0.019] |
|  | Attn head | 0.4564 | 0.4628 | +0.0064 [−0.012, +0.025] |
| Human | Ridge (mean) | 0.5655 | 0.5672 | +0.0017 [−0.013, +0.016] |
|  | Ridge (BOS) | 0.4517 | 0.4686 | +0.0169 [−0.008, +0.042] |
|  | Ridge (summ.) | 0.5502 | 0.5677 | +0.0175 [−0.003, +0.038] |
|  | CNN head | 0.5734 | 0.5977 | **+0.0243 [+0.011, +0.037]** |
|  | Attn head | 0.5711 | 0.5964 | **+0.0253 [+0.011, +0.040]** |
| Human-cell | Ridge (mean) | 0.5472 | 0.5560 | +0.0089 [−0.007, +0.025] |
|  | Ridge (BOS) | 0.4416 | 0.4697 | **+0.0281 [+0.001, +0.054]** |
|  | Ridge (summ.) | 0.5287 | 0.5399 | +0.0112 [−0.012, +0.035] |
|  | CNN head | 0.5463 | 0.5689 | **+0.0226 [+0.009, +0.038]** |
|  | Attn head | 0.5599 | 0.5623 | +0.0024 [−0.015, +0.021] |

On `human_cell`, the CNN head partially replicates the pattern at +0.0226, but the attention head gains only +0.0024, inside seed spread. On the mixed split, no trained head shows a gain, consistent with the phylogenetic structure of that split (Section 6). The mean-pooled ridge gain of +0.0222 does not survive composition control (Section 5.2).

### 5.2. The Gain Survives Composition Control

The improvement on `human` is not explained by amino acid composition, the dominant confound in thermostability prediction. After residualizing the full 400-dimensional dipeptide matrix from both predictions and labels using cross-fitted RidgeCV (Table 2), both trained heads retain clear gains: attention at +0.0521 [+0.025, +0.079] and convolution at +0.0330 [+0.007, +0.059]. The cross-fitting prevents overfitting the residualization itself, and with predictions averaged over 10 seeds inside each of 4000 bootstrap resamples, the intervals reflect protein-sampling uncertainty with readout noise averaged out. That two architecturally different heads agree after a 400-dimensional correction narrows the space of explanations considerably.

**Table 2.**
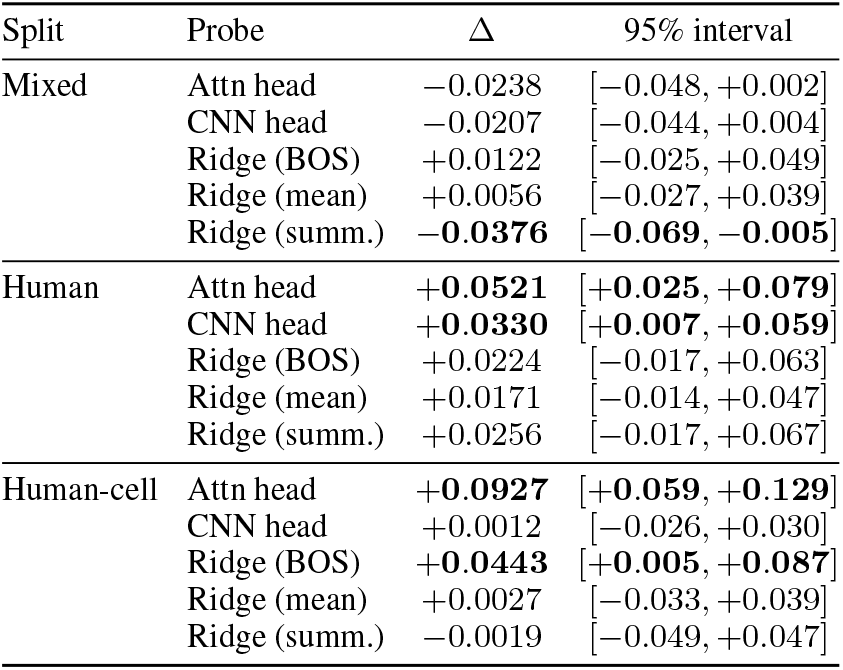
Field-minus-control difference in partial Spearman after residualizing the 400-dimensional dipeptide matrix from predictions and labels (cross-fitted RidgeCV), with 95% paired boot-strap intervals from 4000 resamples of test proteins; trained heads use predictions averaged over 10 seeds inside each resample. Bold marks intervals excluding zero.

| Split | Probe | Δ | 95% interval |
|---|---|---|---|
| Mixed | Attn head | −0.0238 | [−0.048, +0.002] |
|  | CNN head | −0.0207 | [−0.044, +0.004] |
|  | Ridge (BOS) | +0.0122 | [−0.025, +0.049] |
|  | Ridge (mean) | +0.0056 | [−0.027, +0.039] |
|  | Ridge (summ.) | **−0.0376** | **[−0.069, −0.005]** |
| Human | Attn head | **+0.0521** | **[+0.025, +0.079]** |
|  | CNN head | **+0.0330** | **[+0.007, +0.059]** |
|  | Ridge (BOS) | +0.0224 | [−0.017, +0.063] |
|  | Ridge (mean) | +0.0171 | [−0.014, +0.047] |
|  | Ridge (summ.) | +0.0256 | [−0.017, +0.067] |
| Human-cell | Attn head | **+0.0927** | **[+0.059, +0.129]** |
|  | CNN head | +0.0012 | [−0.026, +0.030] |
|  | Ridge (BOS) | **+0.0443** | **[+0.005, +0.087]** |
|  | Ridge (mean) | +0.0027 | [−0.033, +0.039] |
|  | Ridge (summ.) | −0.0019 | [−0.049, +0.047] |

The control split confirms the specificity of the finding: on mixed, no gain survives and one probe (summary-statistic ridge) favors the sequence-only arm at −0.0376 [−0.069, −0.005]. On `human_cell`, the two heads disagree outright (attention at +0.0927, convolution at +0.0012), so we do not count it as a replication.

### 5.3. The Gain Survives Removing Overlapping Proteins

The gain on `human` cannot be attributed to the field arm having seen test proteins’ structures during pretraining. Only the field arm receives structure-derived targets for corpus proteins, so overlap between the corpus and the FLIP test set is a potential confound specific to the ablation. At 0.80 token overlap, 7.0% of `human` test proteins appear in the corpus. Removing all of them barely moves the estimates: the attention head goes from +0.0521 to +0.0518 and the CNN head from +0.0330 to +0.0294, both intervals still clear of zero (Table 14).

This test is conservative: our overlap criterion (at least 0.95 of 3-mers shared with a single corpus protein) is stricter than the one applicable to ESM, where UniProt membership at the clustering identity places at least 91.5–98.3% of Meltome test proteins inside the pretraining set (Appendix K). Our corresponding figure is 2.4–2.9%.

### 5.4. Both Arms Exceed One-Hot and Untrained Baselines

Even at 1.1M parameters, roughly 600 times smaller than ESM-1b, both Biophysical Transformer arms exceed FLIP’s one-hot CNN and untrained-ESM baselines on every split (Table 3). A model this small recovering meaningful thermostability signal suggests that the training objectives, not parameter count alone, are doing useful work.

**Table 3.** FLIP Meltome (Spearman). FLIP rows use FLIP’s own head, optimizer and protocol and are reproduced from the bench-mark, so this is not a controlled comparison. ESM rows are FLIP’s per-AA variants (learned attention pooling, the closest match to our attention head); ESM-1v (Meier et al., 2021) is a single ensemble member. BioTransformer rows use the attention head; italics are composition baselines we add.

| Model | Mixed | Human | Human-cell |
|---|---|---|---|
| ESM-1b (per AA) | 0.68 | 0.71 | 0.76 |
| ESM-1v (per AA) | 0.65 | 0.77 | 0.78 |
| ESM-untrained (per AA) | 0.44 | 0.44 | 0.46 |
| CNN (one-hot) | 0.34 | 0.50 | 0.49 |
| Ridge (one-hot) | 0.17 | 0.15 | 0.24 |
| *AA composition* | *0.3204* | *0.5392* | *0.5438* |
| *Dipeptide (400-d)* | *0.3920* | *0.5121* | *0.5099* |
| **BioTransformer** | **0.4628** | **0.5964** | **0.5623** |
| BioTransformer, seq-only | 0.4564 | 0.5711 | 0.5599 |

The gap to ESM-1b (Rives et al., 2021) remains 0.11–0.22 in Spearman, but the comparison is uncontrolled: FLIP uses a different head, optimizer, protocol and corpus, and untrained ESM provides its head with 1280-dimensional features against our 64, so the absolute numbers are not directly comparable. Only the arm difference is attributable to the field objective. Notably, dipeptide composition alone beats FLIP’s one-hot CNN and ridge on every split, which is why we control for composition directly rather than relying on the baseline comparison.

### 5.5. Transfer to Intrinsic Disorder Prediction (CAID)

To test whether the field signal generalises beyond thermostability, we evaluate frozen BioTransformer embeddings on the CAID disorder-PDB benchmark (319 proteins with DisProt-curated per-residue disorder annotations). A logistic regression probe was trained on per-residue embeddings using 5-fold cross-validation grouped by protein, then scored by the CAID assessment pipeline. We report AUC-ROC and MCC (at a fixed threshold of 0.5). Three embedding sources were compared: BioT voxel (field arm), BioT no-voxel (sequence-only control) and ESM-2 (650M). Published CAID results for AlphaFoldrsa, AIUPred and LIPNet are included as external baselines.

#### Pretraining-set overlap

At 80% sequence identity (Figure 4a), ESM-2 shares similarity with 91% of CAID proteins through its UniRef50 training set. AlphaFold-rsa overlaps with 51% through the PDB. BioT overlaps with just 5%. AIUPred is excluded from the overlap analysis because its deep learning component trains on DisProt, the source of the CAID ground-truth labels.

**Figure 4.**
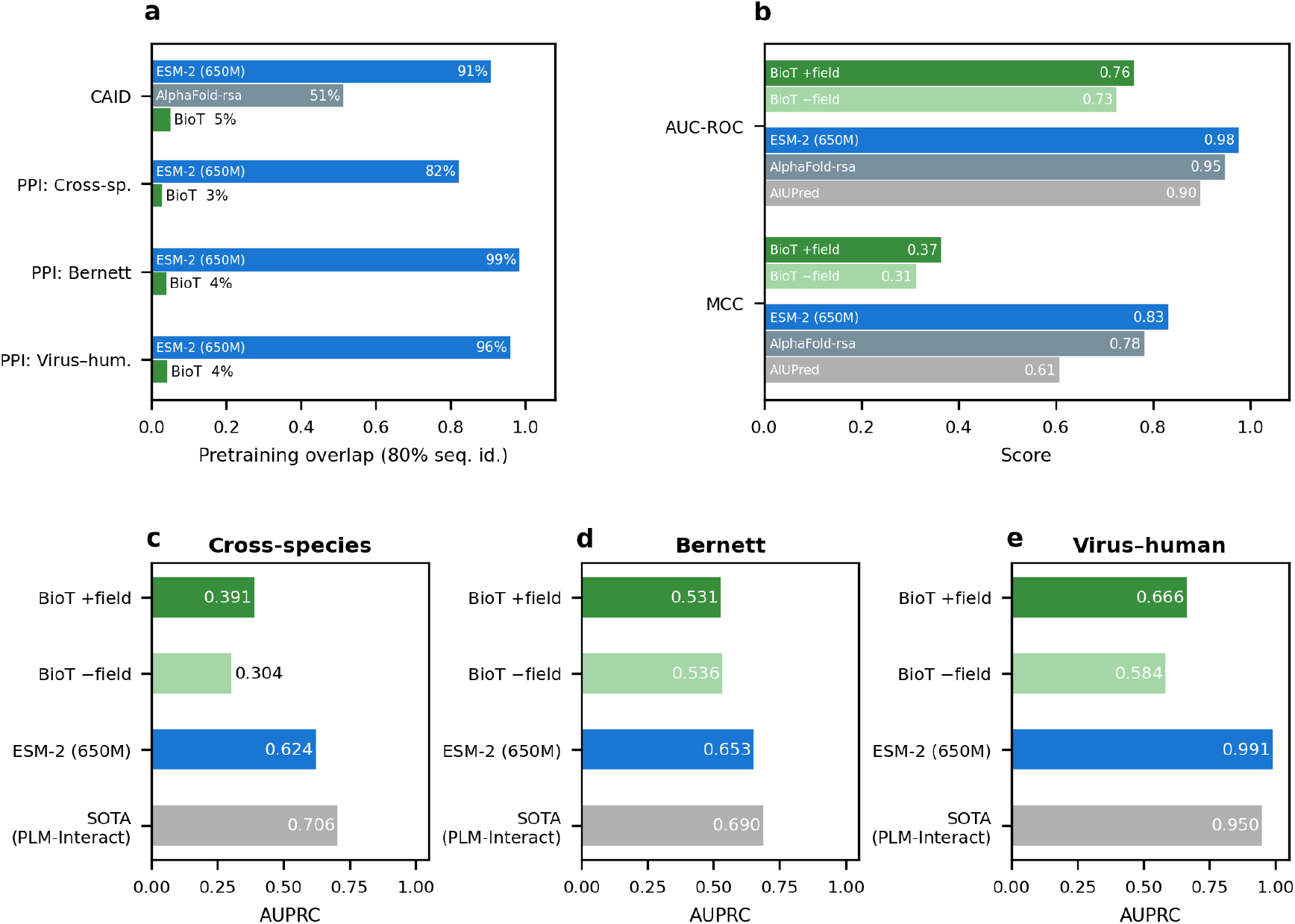
Transfer beyond thermostability. *(a)* Pretraining-set overlap at 80% sequence identity across all transfer benchmarks; AlphaFold overlap (51%) is shown for CAID only, where it is an evaluated baseline. AIUPred trains on DisProt (the CAID label source) and is excluded from the overlap comparison. *(b)* CAID disorder prediction: AUC-ROC and MCC; BioT +field and *−*field denote the field arm and sequence-only control. *(c–e)* PLM-Interact PPI prediction (AUPRC): BioT and ESM-2 use the same frozen-embedding cross-attention probe; SOTA is the published PLM-Interact result with a task-specific architecture.

#### Results

ESM-2 achieves the highest scores (AUC-ROC 0.98, MCC 0.83), followed by AlphaFold-rsa (0.95, 0.78). BioT +field reaches AUC-ROC 0.76 and MCC 0.37; BioT −field reaches 0.73 and 0.31 (Figure 4b). The field objective yields a consistent improvement on both metrics. A paired protein-level bootstrap (2,000 replicates, shared protein resamples across both arms) confirms that the Δ AUC-ROC and Δ MCC distributions lie entirely above zero at the 95% level (Figure 11): the median Δ AUC-ROC is +0.035 and the median Δ MCC is +0.054, with neither distribution approaching the null. Because the same proteins are scored by both models in each replicate, protein difficulty cancels in the paired difference, giving tighter intervals than marginal confidence intervals would suggest.

The absolute gap to ESM-2 must be read alongside two confounds: ESM-2’s corpus covers 91% of the benchmark versus BioT’s 5%, and its embedding dimension (1280) is 20 × larger than BioT’s (64). Despite these disadvantages, BioT +field achieves meaningful per-residue discrimination from a model that has essentially never seen these proteins.

#### Embedding structure

UMAP projections of mean-pooled per-protein embeddings (Figure 10), coloured by ground-truth disorder fraction, reveal qualitative differences in how each model organises the protein space. ESM-2 shows the clearest separation between ordered and disordered proteins, with a smooth gradient along the disorder axis — consistent with its 91% pretraining overlap and 1280-dimensional embedding space. BioT +field produces a more compressed manifold (*d*=64) but still separates highly disordered proteins from ordered ones, with disordered proteins clustering toward distinct regions. Compared to the −field control, the +field model shows tighter grouping by disorder fraction, suggesting that the voxel reconstruction objective imposes structure on the latent space that aligns with biophysical disorder, even for proteins the model has not seen during pretraining. The − field control shows weaker organisation: the disorder gradient is noisier and high-disorder proteins are more diffusely scattered.

### 5.6. Protein–Protein Interaction Prediction

Interaction sites are shaped by the electrostatic complementarity between binding partners, making PPI prediction a downstream task where the field objective may add signal. We evaluate on the PLM-Interact benchmark (Liu et al., 2025), which comprises three tasks: cross-species PPI prediction, Bernett interactome reconstruction and virus–human interaction prediction. For each task, a cross-attention probe was trained over frozen per-residue embeddings; the probe architecture is identical for BioT and ESM-2. Published results for PLM-Interact, TUnA, TT3D, STEP, Topsy-Turvy and D-SCRIPT are included as baselines; all use task-specific architectures and, in most cases, fine-tuned PLM backbones.

#### Pretraining-set overlap

ESM-2’s overlap is extreme (Figure 4a): 82% on cross-species, 99% on Bernett and 96% on virus–human. BioT overlaps with 3–4% across all tasks.

#### Results

Table 4 and Figure 4c–e summarise AUPRC on held-out test sets. BioT voxel achieves 0.391 on cross-species and 0.666 on virus–human. These numbers fall below the published baselines, but the comparison is between a frozen 64-dimensional embedding with a minimal probe and purpose-built PPI predictors using 1280+ dimensional PLMs with task-specific training. The voxel objective improves AUPRC on cross-species (+0.087) and virus–human (+0.082) over the no-voxel control; on Bernett both BioT variants are near chance and the difference is not meaningful.

**Table 4.** PLM-Interact AUPRC. BioT and ESM-2 rows use the same frozen-embedding cross-attention probe; published base-lines use task-specific architectures.

| Method | Cross-sp. | Bernett | Virus–human |
|---|---|---|---|
| PLM-Interact | 0.706 | 0.690 | 0.950 |
| TUnA | 0.641 | 0.690 | — |
| TT3D | 0.553 | — | — |
| STEP | — | 0.620 | 0.893 |
| Topsy-Turvy | 0.490 | 0.580 | — |
| D-SCRIPT | 0.430 | 0.550 | — |
| ESM-2 (probe) | 0.624 | 0.653 | 0.991 |
| **BioT voxel** | **0.391** | 0.531 | **0.666** |
| BioT no-voxel | 0.304 | 0.536 | 0.584 |

### 5.7. MHC Binding Prediction

Peptide–MHC binding affinity depends on the electrostatic complementarity between the peptide and the MHC groove (Wilson et al., 2024), making it a downstream task where the field objective’s spatial information may contribute directly. We compare frozen BioTransformer embeddings against frozen ESM-2 (650M) embeddings on held-out MHC I and MHC II binding sets, using the same regression head (ensemble of 10) for both embedding sources. We report Spearman correlation (SCC), which is threshold-free and therefore robust to differences in prediction calibration between the two models.

BioTransformer outperforms ESM-2 on both tasks (Table 5): Spearman 0.757 versus 0.639 on MHC I (+0.118) and 0.613 versus 0.517 on MHC II (+0.096). This is the one transfer benchmark where the 600× -smaller model exceeds the larger one outright, not merely closes the gap. The result is consistent with the physical argument: MHC binding is governed by charge complementarity in a confined groove, exactly the kind of local electrostatic structure the field objective is trained to reconstruct. The 64-dimensional BioT embedding, shaped by explicit electro-static supervision, apparently encodes binding-relevant information that ESM-2’s 1280-dimensional sequence-only embedding does not capture as efficiently.

**Table 5.**
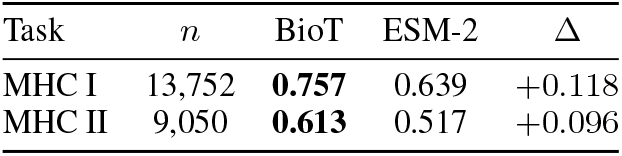
MHC binding prediction (Spearman). Both models use the same frozen-embedding regression head (ensemble of 10). BioT is the field arm.

| Task | *n* | BioT | ESM-2 | Δ |
|---|---|---|---|---|
| MHC I | 13,752 | **0.757** | 0.639 | +0.118 |
| MHC II | 9,050 | **0.613** | 0.517 | +0.096 |

## 6. Discussion

### 6.1. Interpretation

The effect appears on the human splits and not on the mixed split. Two explanations predict exactly this pattern. First, between-organism variance may be largely phylogenetic and already captured by MLM, leaving little for the field objective to add when organisms are pooled. Stratifying the mixed split by organism (Appendix M) supports the premise: the control’s Spearman falls from 0.456 on the whole split to between 0.079 and 0.264 within single organisms, so most of the mixed split’s predictability lies between organisms rather than within them. Second, the corpus’s human enrichment (18.1% against 3.1% for *Mus musculus*) may have given the field objective more human proteins to learn from. The stratification cannot adjudicate between these two causes: no single organism shows a separable difference, and the test lacks the power to detect one.

On `human_cell`, the two trained heads disagree outright after composition control (attention +0.0927, convolution +0.0012), and the attention figure is unstable under protein removal (Table 14). We do not count this split as a replication.

More fundamentally, the control tests whether the field objective changes the encoder, but it cannot separate two mechanisms. The improvement may come from the physical content of the field (continuum electrostatics, solvation parameters) or from the regularization effect of any auxiliary objective at matched gradient scale. METL (Gelman et al., 2025) faces the same question in a different geometry: its Rosetta energy targets are global scalars, so a permuted-energy control is straightforward, but it has not been reported. In our setting, a permuted-target arm would separate these explanations, and an atom-density target with the same descriptor would directly test the physical-prior hypothesis. Neither has been run. The field is a deterministic function of the coordinates, and what makes it distinctive is not continuity or frame-freeness (structure can be supervised through frame-free targets) but the physical model’s own assumptions, its fixed parameters and solvation model, none of which sit in a coordinate list. Whether those assumptions act as a useful prior is a question of inductive bias that is not resolved here.

### 6.2. Limitations

The claim rests mainly on `human`, and a human-specific effect has the two candidate causes above. The field targets come from preprocessing code and solver settings that are not released, so the pretraining corpus cannot be regenerated and the pretraining stage is not reproducible from this report. Only one of seven degree blocks supervises polarity: the *l*=0 shell means are linear in the field, but all angular terms are quadratic and therefore sign-blind. Negating a perfect prediction would raise the descriptor distance from 0 to 2*/*7, all of it in the *l*=0 block (Appendix E). Appendices D and F detail additional technical limitations: the centered distance does not constrain relative orientation between residues, the frame assumption is approximate, and the tokenization parameter *k*=3 was not varied.

## 7. Future Work

Three controls would sharpen attribution: a second pre-training run per arm, to separate encoder variance from the field effect; an equally invariant atom-density target, the direct test of the physical-prior argument; and a permuted-target arm at matched gradient scale, separating content from regularization. Beyond these controls, the most immediate extensions are a scale study over *k* (with hashed or factorized embeddings for *k* ≥ 4), the objective at ESM scale to test whether it stays additive, and a decoder that represents a frame shared across residues so that relative arrangement can be supervised.

### Electrostatics beyond biology

The Poisson–Boltzmann field is not a biological concept; it is a solution to a PDE on a charge distribution in a dielectric medium. Nothing in the descriptor or the training objective is specific to amino acids. In principle the same framework could pre-train representations for any molecular system that generates a charge-driven field: small-molecule solvation, polymer electrolytes, or nucleic acid complexes. Whether the invariant descriptor learned here transfers to non-biological electrostatics is untested, but the architecture imposes no domain restriction.

### From reconstructed fields to structure

The decoder already produces a 33^3^ potential grid per token. These grids are low-fidelity at the current corpus size, but they are genuine volumetric predictions conditioned only on sequence. With a larger training corpus and a higher-resolution decoder, the predicted fields could in principle be inverted to recover the charge distribution and hence the atomic positions. This route to structure prediction would avoid the *N* ^2^ distance-matrix representation that limits many current approaches: each token’s field depends primarily on its local covalent neighbours, so the representation scales linearly with sequence length even though the physics it captures is three-dimensional. The preliminary CAID and PPI results (Sections 5.5 and 5.6) already suggest that the encoder learns something about local spatial arrangement.

### Condition-dependent fields

The current corpus uses a single set of APBS parameters (PARSE charges, 150 mM ionic strength, pH 7). The electrostatic field of a residue changes with solvent conditions: pH shifts protonation states, ionic strength screens long-range interactions, and divalent cations create specific binding geometries. A natural extension is to condition the decoder on these parameters, so that the same encoder produces condition-dependent predictions. The same three-residue token already appears in different protein environments across the corpus; extending to salinity, pH and ion type would make the model predict how a peptide’s field changes with its physical context, not only its sequence context.

## 8. Conclusion

Figure 3 summarizes the Biophysical Transformer’s contribution along three axes. At ×600 fewer parameters and 4,700× fewer pretraining sequences than ESM-1v, the field arm reaches a Spearman *ρ* of 0.60 on the `human` thermostability split, closing nearly half the gap from an untrained 650M-parameter encoder. This performance cannot be attributed to benchmark exposure: only 7% of test proteins appear in the pretraining corpus, versus ∼95% for ESM-1v, and removing all overlapping proteins barely moves the estimate. The raw field-minus-control difference of +0.025 accounts for 8% of the untrained-to-ESM-1v gap; after residualizing dipeptide composition, the advantage grows to +0.052, with both the attention and CNN heads’ 95% intervals clear of zero. Nothing survives on the mixed split, and the current design cannot separate field content from regularization, but the result establishes the electrostatic potential as a usable auxiliary pretraining signal for protein language models. Transfer experiments on intrinsic disorder (Section 5.5), protein–protein interaction (Section 5.6) and MHC binding (Section 5.7) confirm that the field signal encodes information beyond thermostability. On MHC I and MHC II, the 600× -smaller Bio-Transformer outperforms ESM-2 outright, consistent with the electrostatic nature of peptide–groove complementarity. These transfer experiments lack the controlled ablation of the Meltome study, but the consistency of the field arm’s advantage across four distinct tasks narrows the space of explanations. The next experiment is the Biophysical Transformer objective at ESM scale.

## Code and Data Availability

Code is available at https://github.com/patjiang/Biophysical-Transformer. The downstream benchmarks (FLIP Meltome (Dallago et al., 2021), CAID, PLM-Interact (Liu et al., 2025)) are public. The pretraining corpus (21K electrostatic potential grids computed by APBS on CATH structures) is being prepared for public release; a hosting location has not yet been finalised.

### Impact Statement

This work studies a pretraining signal for protein representation learning. Such representations carry the field’s usual dual-use character, but the model is small, trained on public structures and evaluated on one public benchmark, so it confers no capability beyond released PLMs. Field targets come from published solvation software on deposited structures and involve no human-subject data.

## Supplementary Material

### A. Objective Weights

Table 6 gives the trained objective in full; every row is read from the epoch-75 checkpoint.

**Table 6.** Trained objective weights (Appendix E). The latent collapse guard is inactive in the control; the encoder guard runs in both.

| Term | Weight |
|---|---|
| Field reconstruction (outer weight) | 1.00 |
| residue-centered descriptor | 1.00 |
| uncentered descriptor | 0.50 |
| radial energy | 0.50 |
| excess total variation | 0.25 |
| all voxel-wise terms | 0.00 |
| cross-residue (both variants) | 0.00 |
| Masked field reconstruction (effective) | 3.00 |
| MLM | 2.00 |
| DINO | 0.10 |
| iBOT | 1.00 |
| Encoder collapse guard | 0.10 |
| Latent collapse guard | 0.10 |

### B. Hyperparameters

MLM uses a span mask of 3 residues at rate 0.2, replacement fraction 0.2 and copy fraction 0.0. Masking is at residue level: with stride-1 *k*-mers each anchor hides *W* +*k* − 1=5 tokens, exactly those containing any of its three residues, with anchor probability 1 −(1 −*ρ*)^1*/*(*W* +*k−*1)^, so a masked residue is absent from the input altogether and the masked-position comparison measures inference from context, not tokenization leakage.

DINO uses 128 prototypes, a 4096-entry queue, teacher momentum 0.999 and a 60,000-step warmup (59% of steps to epoch 75). iBOT uses 256 prototypes, an 8192-entry queue, temperatures 0.1 (student) and 0.05 (teacher), and a teacher band of 2 residues per layer, which over the encoder’s 5 layers is a receptive field of ±10 residues.

The collapse guards act on three statistics of the states *z*, each penalized only below its target so that the term vanishes once the target is met. Let *C* be the covariance of the states centered within protein. *Scale* is their root mean square, penalized as max(0, 1.0 − rms). *Participation ratio* is 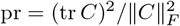, equal to 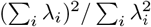 over the eigenvalues of *C*, so it counts the directions carrying variance: 1 if one direction carries all of it, *d* if variance is spread evenly over *d*. Against a target pr^∗^=20.0 of the available 64, the penalty is max(0, log(pr^∗^*/*pr)*/* log pr^∗^). *Share* is the fraction of the states’ raw mean square carried by the within-protein component, 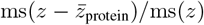, penalized as max(0, 0.25 −share)*/*0.25; it exists because the first two statistics can be satisfied by variation between proteins while the states are nearly constant along each one.

The vocabulary has 8003 entries (8000 3-mers plus `PAD`, `MASK`, `BOS`). The interaction block uses layer scale 0.3 and token initialization 0.1; the identity skip is a separate module whose gate starts at exactly 0.0. The MLM head carries a unigram bias scaled by 0.25, giving an expected initial cross-entropy of 8.8353.

### C. Decoder Architecture

The decoder (Figure 5) refines the encoder’s per-token output *z* with a residual block of 32 learned interaction tokens (2 heads) and a zero-gated skip connection from the input embeddings. Learned, non-equivariant projections of the BOS state and per-token features build a 29-channel seed volume on a fixed 4^3^ coordinate grid, which fixes the output frame. One SE(3)-equivariant cross-attention layer (4^3^) and one equivariant transformer layer (8^3^) refine it, and a convolutional path with FiLM conditioning upsamples via trilinear interpolation to 16^3^ then 33^3^, with 1 × 1 channel reductions at each stage. The output is closed by a tanh nonlinearity (field bounded to ± 4981.9 before normalization). Of 1.1M trainable parameters, 398K sit in the decoder path: 341K in the voxelization trunk, 52K in the interaction-token block, and 4K in the identity projection. We did not ablate the equivariant layers.

**Figure 5.**
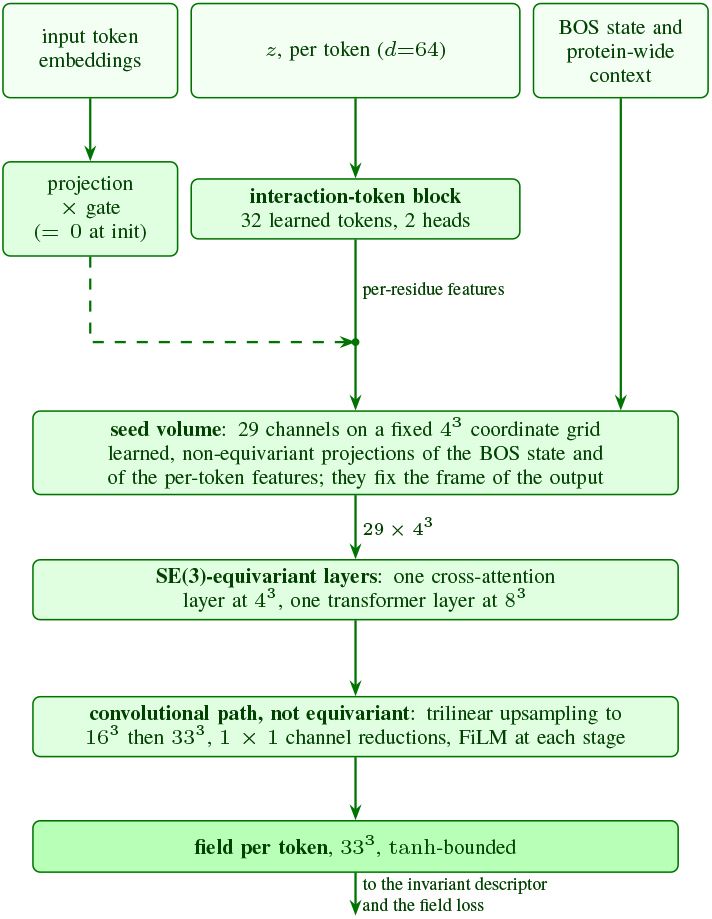
The field decoder. (field arm only, green). Inputs come from the shared encoder. Non-equivariant projections build the seed volume and fix the output frame; two equivariant layers act at 4^3^ and 8^3^ and the rest is convolutional. The invariant descriptor scores the result (Figure 1).

**Figure 6.**
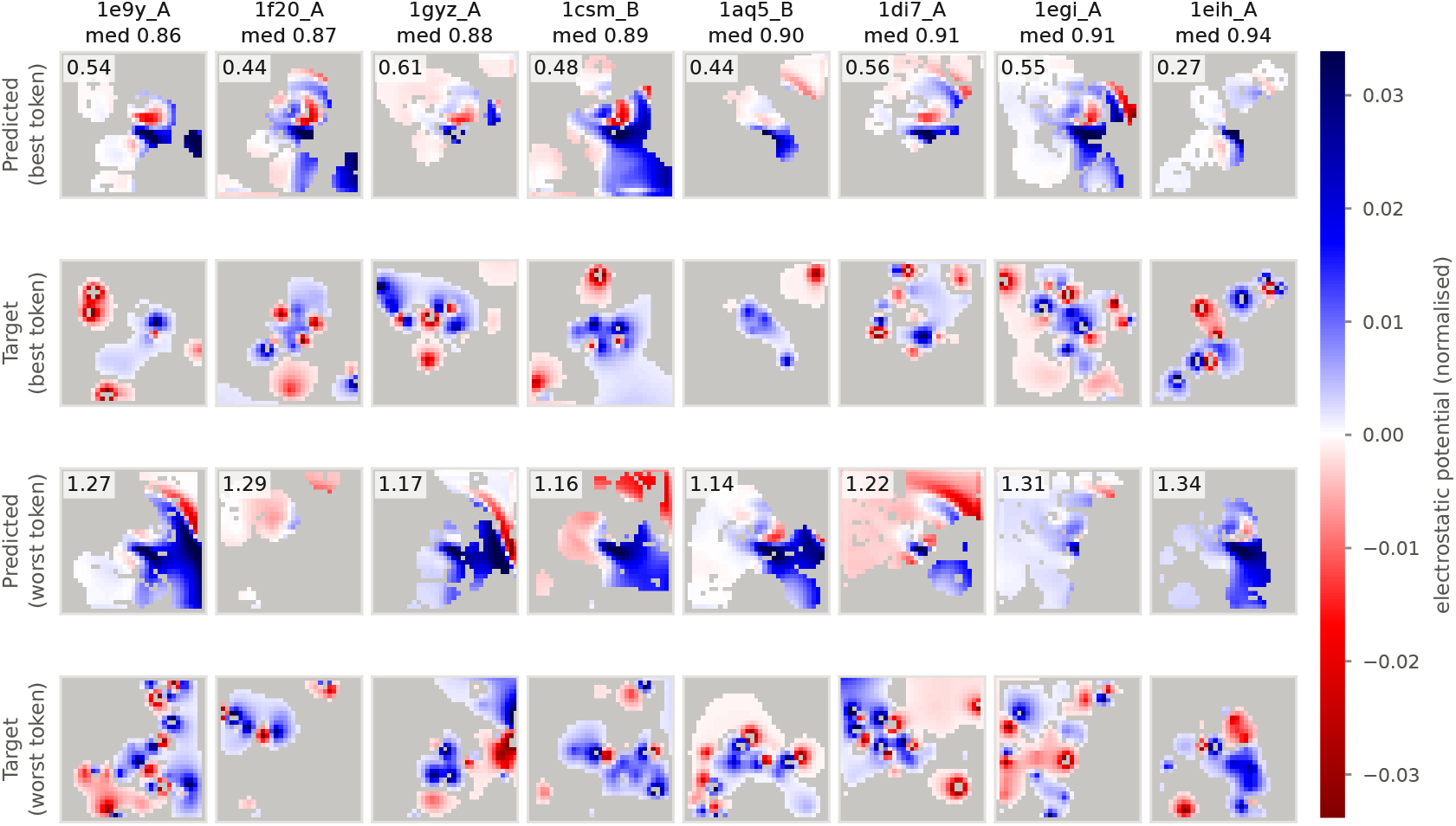
Electrostatic potential volumes for representative proteins. Mid-plane slices of the 33^3^ box for eight held-out proteins, ordered left to right by median descriptor error at evenly spaced percentiles from the 5th (1e9y_A, 0.856) to the 95th (1eih_A, 0.940). For each protein the best and worst tokens are shown, bracketing the within-protein range; column headers give the protein’s median error and each predicted panel carries that token’s own. Panels show the raw field, not the within-protein residual, so a component common to a protein’s tokens is visible here but enters none of the errors in Table 11. All panels share one colour scale, symmetric about zero and clipped at the 99.5th percentile of |value| . Positions whose 3-mer window runs off a chain break are excluded. Because the objective is rotation-invariant, orientation is never supervised and the predicted and target panels are not expected to align.

### D. The Invariant Descriptor

The 33^3^ field is summarized on 12 Gaussian shells (*σ*=1.0 voxel), centered from *r*=2.0 to 16.0 voxels (the outermost at a face-voxel center), quadrature-normalized. Shells start at *r*=2 because one at *r* ≈1 holds about a dozen voxels and cannot resolve *l*=6. Coefficients up to *l*_max_=6 give 49 per shell, 588 in all. The descriptor concatenates the 12 *l*=0 values with the 12 × 12 cross-shell matrices *P*_*l*_(*s, s*^*′*^) = Σ _*m*_ *c*_*lm*_(*s*)*c*_*lm*_(*s*^*′*^) for *l*=1, …, 6, for 12 + 6 × 12^2^ = 876 dimensions; coefficients are not *L*^2^-normalized, and *l* ≥ 1 blocks are compressed as sign(*P* )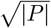 .

Equation (2) is applied *per position* and the values then averaged, not to a protein’s stack of descriptors as a whole: the inner product and both norms run over the descriptor dimension of one position, one degree block at a time, and the resulting distances are averaged first over the protein’s positions and then over the seven blocks. A protein therefore contributes one value per block regardless of its length.

The descriptor is exactly invariant under the 24 grid rotations (distance 0.000000). Under random continuous rotation, which needs resampling, the median distance is 0.0136 and the 90th percentile 0.0177 (25 proteins), a lattice effect. That average conceals a hundredfold spread: measured per degree block, the error rises from 0.0005 at *l*=0 to 0.0501 at *l*=6 (Table 7), since a fixed lattice resolves high angular degrees least well. The decoder’s margin over the strongest baseline at each degree exceeds the rotation error by a factor of 40 at *l*=0, 11 at *l*=2, 5 at *l*=3, but only 1.5 at *l*=5 and less than one at *l*=6. The per-degree advantage is therefore established up to *l*=4 and unresolved at the two highest degrees.

**Table 7.** Descriptor distance by degree block, on the same 400 proteins as Table 11; lower is better. Bold marks the lowest value where the model’s margin over the next-best predictor is at least twice the rotation error.

| Block | Residue-type lookup | Window ridge, ±5 | 3-mer lookup | Model | Rot. err. |
|---|---|---|---|---|---|
| *l*=0 | 0.938 | 0.809 | 0.840 | **0.788** | 0.0005 |
| *l*=1 | 0.984 | 0.950 | 0.944 | **0.907** | 0.0019 |
| *l*=2 | 0.984 | 0.948 | 0.939 | **0.896** | 0.0039 |
| *l*=3 | 0.983 | 0.953 | 0.944 | **0.909** | 0.0067 |
| *l*=4 | 0.984 | 0.957 | 0.948 | **0.912** | 0.0137 |
| *l*=5 | 0.987 | 0.962 | 0.956 | 0.919 | 0.0238 |
| *l*=6 | 0.989 | 0.965 | 0.962 | 0.925 | 0.0501 |
| Mean, *l* ≥ 1 | 0.985 | 0.956 | 0.949 | **0.911** | 0.0167 |
| All blocks | 0.9784 | 0.9349 | 0.9335 | **0.8937** | 0.0147 |

The loss mask (*r* ≤ 16.5) keeps 18,853 of 35,937 voxels.

### E. Loss Terms

The residue-centered descriptor agreement term (weight 1.0) is the primary loss: it computes *d* after subtracting each protein’s mean descriptor, so it measures residue-to-residue variation. The uncentered agreement term (0.5) pins the protein-wide level. The radial energy term (0.5) matches the shell mean of *V* ^2^ and is the only term accounting for energy above *l*=6. The excess total variation term (0.25) penalizes striping that reproduces a power spectrum at higher gradient than the target. All voxel-wise terms have zero weight.

A cross-residue term, comparing spherical-harmonic inner products for residue pairs within 10 positions, is implemented but did not run: the outer weight of the block it sits in is zero throughout training. The four active terms are each computed independently at each residue, so they are invariant to a separate rotation of every residue’s field. This is a larger group than Equation (1) licenses.

Masked field reconstruction applies the same objective at masked positions with effective weight 3.0, the largest in the objective. There the decoder receives the mask embedding and the residue is absent from the encoder input, so the loss falls only insofar as *z* carries the hidden residue’s environment from context.

Only the *l*=0 block distinguishes *V* from −*V* : negating a perfect prediction raises both descriptor-agreement terms from 0 to 2*/*7 ≈ 0.286, all in that block. This follows from the construction, not from any checkpoint.

### F. The Frame Assumption

Equation (1) rests on two conditions: that each box is axis-aligned to the deposited structure’s frame (holds by construction), and that the molecule’s orientation within that frame is uniform over the 24 grid-preserving rotations *G* and independent of the sequence.

Uniformity over *G* makes E[*V* ] *G*-invariant, which for *l* ≤ 6 requires no power at *l* = 1, 2, 3, 5. Over 2,000 held-out proteins against 1,000 null realizations that impose uniformity by construction (Table 8), the dipole sits at 2.7 times the null median (*p* = 0.13 after Holm correction across the four tested degrees); the other three are at 0.82 or above.

**Table 8.** Spherical-harmonic power of the corpus-mean target field, by degree, against a null imposing uniformity. 2,000 held-out proteins, 1,000 null realizations. The ratio is observed over null median. *p*_Holm_ corrects across the four degrees at which the group admits no invariant; the other three check that the null behaves.

| *l* | Observed | Null median | Ratio | *p*<sub>Holm</sub> |
|---|---|---|---|---|
| 0 | 3.50 × 10<sup>−4</sup> | 3.50 × 10<sup>−4</sup> | 1.00 | — |
| 1 | 1.47 × 10<sup>−8</sup> | 5.48 × 10<sup>−9</sup> | 2.69 | 0.128 |
| 2 | 6.32 × 10<sup>−9</sup> | 6.84 × 10<sup>−9</sup> | 0.92 | 1.000 |
| 3 | 3.34 × 10<sup>−9</sup> | 2.83 × 10<sup>−9</sup> | 1.18 | 0.821 |
| 5 | 1.38 × 10<sup>−9</sup> | 1.56 × 10<sup>−9</sup> | 0.88 | 1.000 |
| 4 | 1.19 × 10<sup>−7</sup> | 1.18 × 10<sup>−7</sup> | 1.00 | — |
| 6 | 6.23 × 10<sup>−8</sup> | 6.25 × 10<sup>−8</sup> | 1.00 | — |

Subsampling at *n* = 250, 500, 1000, 2000 (Table 9), the dipole power falls as *n*^*−*0.61^ while the other tested degrees fall as *n*^*−*0.99^ to *n*^*−*1.16^, and every null falls as *n*^*−*1.03^ to *n*^*−*1.08^. Uniformity therefore fails at the dipole alone.

**Table 9.**
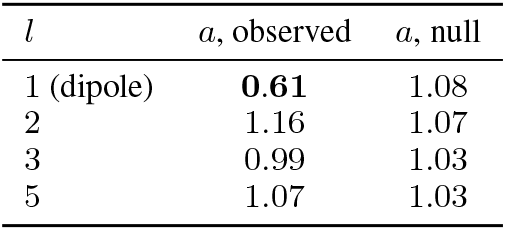
Decay exponent *a* of degree power with sample size, power ∝ *n*^*−a*^, over *n* = 250 to 2,000. Uniformity requires *a* = 1.

| *l* | *a*, observed | *a*, null |
|---|---|---|
| 1 (dipole) | **0.61** | 1.08 |
| 2 | 1.16 | 1.07 |
| 3 | 0.99 | 1.03 |
| 5 | 1.07 | 1.03 |

The coherent dipole is 3.2% of a typical protein’s own dipole amplitude and 2.9 ×10^*−*5^ of the *l* = 0 power. Since it is a single direction shared by the corpus and identical for every S, it distinguishes no residue from another and the collapse argument of Section 3.3 survives in kind.

A translation of the boxes is excluded by the measured support centroid (mean displacement 0.0096 voxels, s.e. 0.0038 per axis), which leaves a mildly non-uniform deposited orientation distribution as the surviving cause. Independence of orientation from the sequence is untested.

### G. Ablation Checks

The two arms are meant to differ in one flag. Algorithm 1 is the procedure that tests this, with the observed value of each assertion beside it. All six checks pass and none fails; the script is `verify_ablation.py` in the released code.

#### Algorithm 1

Ablation fairness checks, seed 0. Each line asserts; the comment gives the observed quantity.

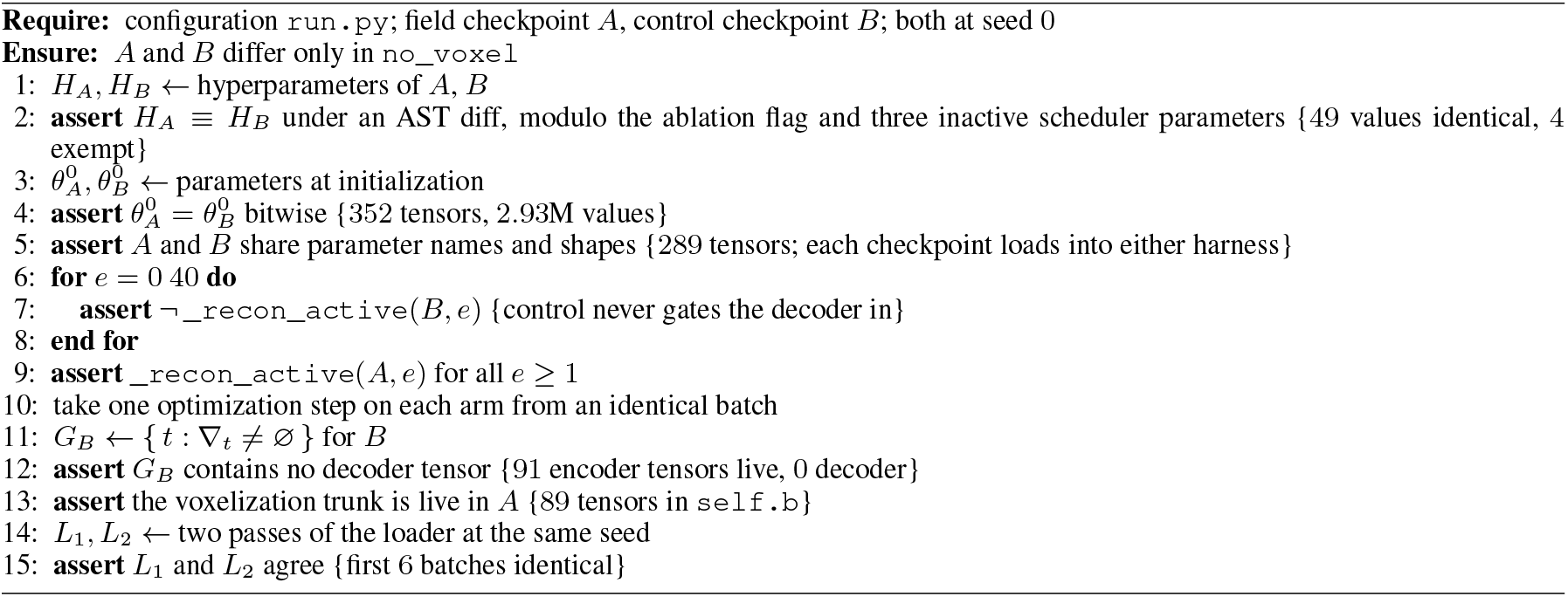

The fifth and sixth checks are the data-backed ones: they run the model rather than compare files, which is what distinguishes a decoder that is configured off from one that is absent from the gradient graph.

### H. Protocol

Heads use the 10 paired seeds 0 to 9. Table 1 reports the mean over seeds of each seed’s Spearman; Table 2 uses each arm’s predictions averaged over seeds inside each bootstrap resample, so its intervals are protein-sampling uncertainty with head noise averaged out. We never compare arms on single seeds, because checkpoint selection noise is applied independently per arm and does not cancel. Single-seed predictions gave anomalous results on two split–head combinations (−0.0895 for the CNN head on `human_cell` and −0.0410 for attention on mixed); under 10-seed averaging both cross zero.

Each arm’s encoder is taken at its best validation loss (epoch 75 field, 79 control). As a sensitivity check, evaluating both at epoch 79 (deterministic probes only) moves composition-controlled deltas by at most 0.0028 on the human splits (0.0128 on mixed, where composition dominates and the residual is small).

### I. Trajectory

The mean-pooled effect decays after epoch 34 while the BOS-pooled effect rises and plateaus (Table 10), so a single-checkpoint result depends on the selection rule (Appendix H). The skip gate reaches 1.55 by epoch 75 (projection weight norm 34.8), so the token pathway is open. The log’s residue-centered distance is flat over the last ten epochs (0.8708– 0.8734); these are tail epochs of the cosine schedule (learning rate under a tenth of peak), so flatness does not indicate convergence.

**Table 10.** Composition-controlled delta across training, both arms at the same epoch.

| Split | e34 | e69 | e79 |
|---|---|---|---|
| *Ridge, mean-pooled* |  |  |  |
| Mixed | +0.0061 | −0.0222 | −0.0072 |
| Human | +0.0621 | +0.0271 | +0.0199 |
| Human-cell | +0.0523 | +0.0039 | +0.0028 |
| *Ridge, BOS* |  |  |  |
| Mixed | +0.0057 | +0.0126 | +0.0198 |
| Human | −0.0145 | +0.0294 | +0.0248 |
| Human-cell | +0.0087 | +0.0469 | +0.0471 |

### J. Field Reconstruction

The downstream result is interpretable only if the decoder learned more than a lookup. Table 11 compares the decoder’s descriptor distance (Equation (2), per block after residue-centering, averaged; lower is better) with three sequence-only predictors: a residue-type lookup, a 3-mer lookup keyed on the encoder’s own token, and a ridge regression on a ±5 residue window. They are fitted on 1,500 training proteins and all rows are scored on the same 400 held-out proteins; refitting on 21,418 proteins moves them by at most 0.0006. A prediction that does not vary across residues scores 1.0, and one with exact shell means and no angular structure scores 6*/*7 ≈ 0.857.

**Table 11.** Held-out descriptor distance (Equation (2)), residue-centered, averaged over seven blocks; lower is better. Rows above the rule use the same 400 held-out proteins; s.e. is between-protein only. Rows below are scored at masked positions, centered among themselves, so comparable with each other but not with rows above.

| Predictor | Distance ↓ | s.e. |
|---|---|---|
| Corpus-mean field | 1.0000 | — |
| Residue-type lookup | 0.9784 | 0.0003 |
| Sequence window, ±5 (ridge) | 0.9349 | 0.0008 |
| 3-mer lookup | 0.9335 | 0.0010 |
| **Model, unmasked input** | **0.8937** | **0.0013** |
| Reference: exact shell means, no angular structure | 6/7 ≈ 0.857 | — |
| *Masked positions, centered among themselves* |  |  |
| Model, residue visible | 0.8838 | — |
| Model, residue hidden | 0.9258 | — |

The decoder reaches 0.8937, against 0.9335 for the 3-mer lookup and 0.9349 for the window ridge. Its advantage is resolvable at degrees up to *l*=4 given the descriptor’s own rotation error (Table 7), and its angular blocks supply 71.5% of its margin below the floor. With the residue masked and absent from the input, the distance rises to 0.9258 against 0.8838 visible, so the decoder keeps about two-thirds of its margin from context alone, as the masked term demands.

The held-out proteins also selected the checkpoint, and 4.5% share a token set with a training protein. Among the 400 scored proteins, 32 are duplicates; the margin over the 3-mer lookup is +0.0366 on those and +0.0394 on the other 368, so duplicates do not carry it. This is evidence the objective was met, not a generalization estimate.

### K. Exposure Audit

A gain measured on held-out proteins means nothing if those proteins were in the pretraining corpus. For each FLIP test protein we compute the fraction of its distinct 3-residue words present in any single pretraining protein (a maximum over the 26,773 corpus proteins, so a test protein whose 3-mers can be assembled from fragments of a thousand corpus entries is not flagged). We calibrate by shuffling each test sequence five times, which preserves length and composition and isolates the part of the overlap that is sequence identity rather than compositional chance; proteins with fewer than 30 distinct words are excluded. No shuffled sequence in any split reaches 0.80, so that threshold marks the boundary of what composition alone can produce.

At 0.80, 7.0%, 8.5% and 5.8% of the scored test proteins are essentially wholly in one corpus protein (Figure 7, left; Table 12). The bulk of each split is clean: medians (0.354–0.357) sit only 0.015–0.020 above shuffled (0.334–0.342), while the 99th percentile jumps to 0.988 or above against a shuffled 99th below 0.50. Exposure is therefore a small contaminated tail on an otherwise clean distribution, precisely the structure that removal can address. Checking the reverse direction confirms that nearly all flagged proteins are whole-protein duplicates, not fragments of longer corpus entries (Table 13): only 1, 1 and 3 fragments across the three splits.

**Table 12.**
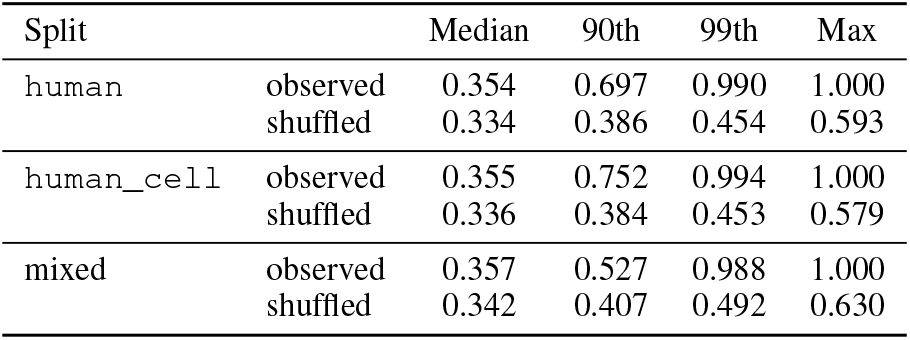
Overlap with the best-matching pretraining protein, observed and for sequences shuffled five times.

| Split |  | Median | 90th | 99th | Max |
|---|---|---|---|---|---|
| human | observed | 0.354 | 0.697 | 0.990 | 1.000 |
|  | shuffled | 0.334 | 0.386 | 0.454 | 0.593 |
| human_cell | observed | 0.355 | 0.752 | 0.994 | 1.000 |
|  | shuffled | 0.336 | 0.384 | 0.453 | 0.579 |
| mixed | observed | 0.357 | 0.527 | 0.988 | 1.000 |
|  | shuffled | 0.342 | 0.407 | 0.492 | 0.630 |

**Table 13.**
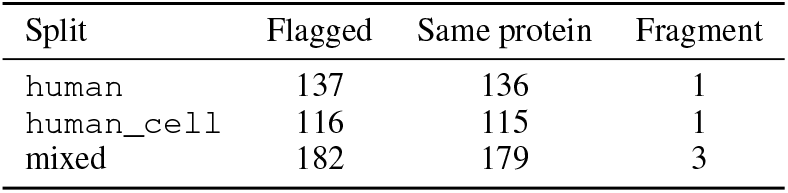
Test proteins flagged at 0.80, by whether the match is the same protein or the test protein is a fragment of a longer entry.

**Table 14.** Composition-controlled field-minus-control difference (partial Spearman) on the full test set and with proteins overlapping the pretraining corpus at 0.80 removed. The full-set column reproduces Table 2. Bold marks an interval excluding zero.

| Probe | Full set | Overlap removed |
|---|---|---|
| human *n* = 1945 → 1808 |  |  |
| Attention | **+0.0521 [+0.025, +0.079]** | **+0.0518 [+0.022, +0.080]** |
| CNN | **+0.0330 [+0.007, +0.059]** | **+0.0294 [+0.002, +0.056]** |
| Ridge (BOS) | +0.0224 [−0.017, +0.063] | −0.0033 [−0.047, +0.039] |
| Ridge (mean) | +0.0171 [−0.014, +0.047] | +0.0124 [−0.020, +0.045] |
| Ridge (summ.) | +0.0256 [−0.017, +0.067] | +0.0269 [−0.015, +0.070] |
| human_cell *n* = 1366 → 1250 |  |  |
| Attention | **+0.0927 [+0.059, +0.129]** | **+0.1744 [+0.133, +0.220]** |
| CNN | +0.0012 [−0.026, +0.030] | −0.0061 [−0.034, +0.022] |
| Ridge (BOS) | **+0.0443 [+0.005, +0.087]** | +0.0271 [−0.017, +0.074] |
| Ridge (mean) | +0.0027 [−0.033, +0.039] | +0.0195 [−0.019, +0.059] |
| Ridge (summ.) | −0.0019 [−0.049, +0.047] | +0.0156 [−0.032, +0.061] |
| mixed *n* = 3134 → 2952 |  |  |
| Attention | −0.0238 [−0.048, +0.002] | −0.0252 [−0.052, +0.000] |
| CNN | −0.0207 [−0.044, +0.004] | −0.0148 [−0.038, +0.010] |
| Ridge (BOS) | +0.0122 [−0.025, +0.049] | +0.0122 [−0.025, +0.050] |
| Ridge (mean) | +0.0056 [−0.027, +0.039] | −0.0096 [−0.042, +0.023] |
| Ridge (summ.) | **−0.0376 [−0.069, −0.005]** | **−0.0333 [−0.066, −0.001]** |

**Figure 7.**
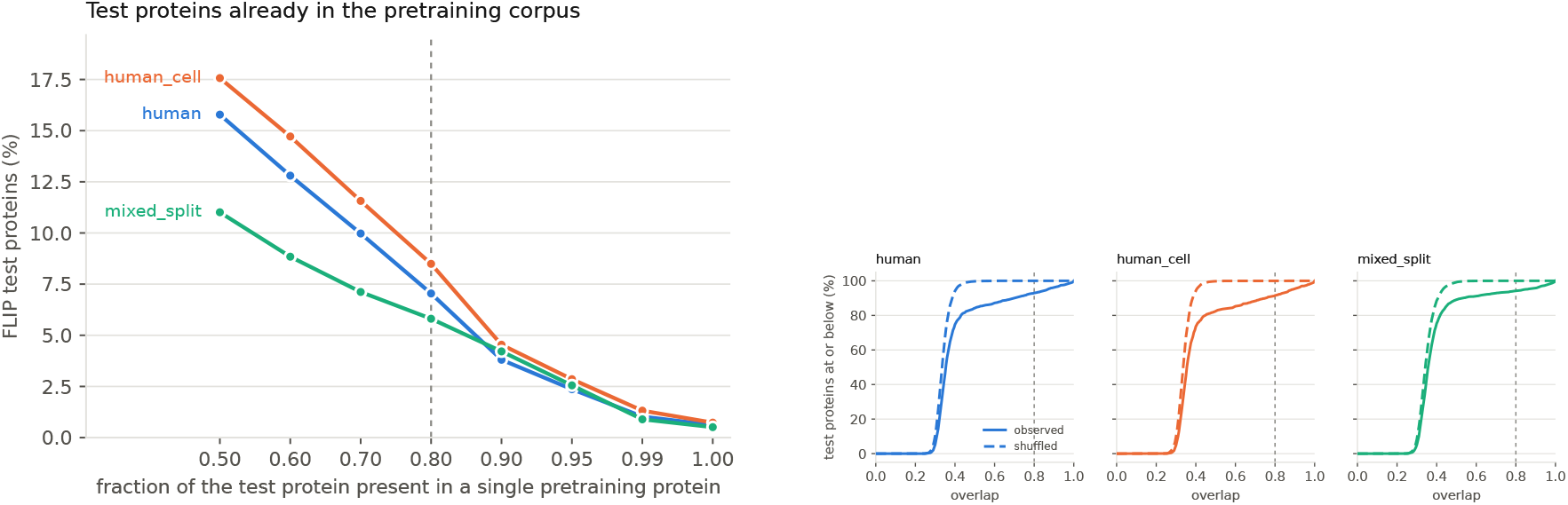
**Left:** Percentage of each split’s test proteins with at least the stated fraction of 3-mers in a single pretraining protein; dashed line marks 0.80. **Right:** Cumulative distribution of per-protein overlap for test proteins (solid) and their shuffles (dashed).

The two criteria in Figure 8 are not the same measurement. Ours asks whether a specific sequence is almost wholly inside one of 26,773 proteins we can enumerate. The ESM figure asks whether a test protein is a member of UniProt at the identity its corpus was clustered to, a far weaker condition satisfied by any protein with a database entry. ESM-1v used UniRef90 release 2020-03 (98M sequences) and ESM-1b UniRef50 release 2018-03 (≈ 30M sequences); UniProt matched 98.3%, 98.4% and 92.7% of test proteins as lower bounds (Figure 8, left). The asymmetry is the point: we apply the stricter criterion to the Biophysical Transformer and the looser one to the baselines, so the gap between the bars understates rather than overstates how much more exposed the baselines are.

**Figure 8.**
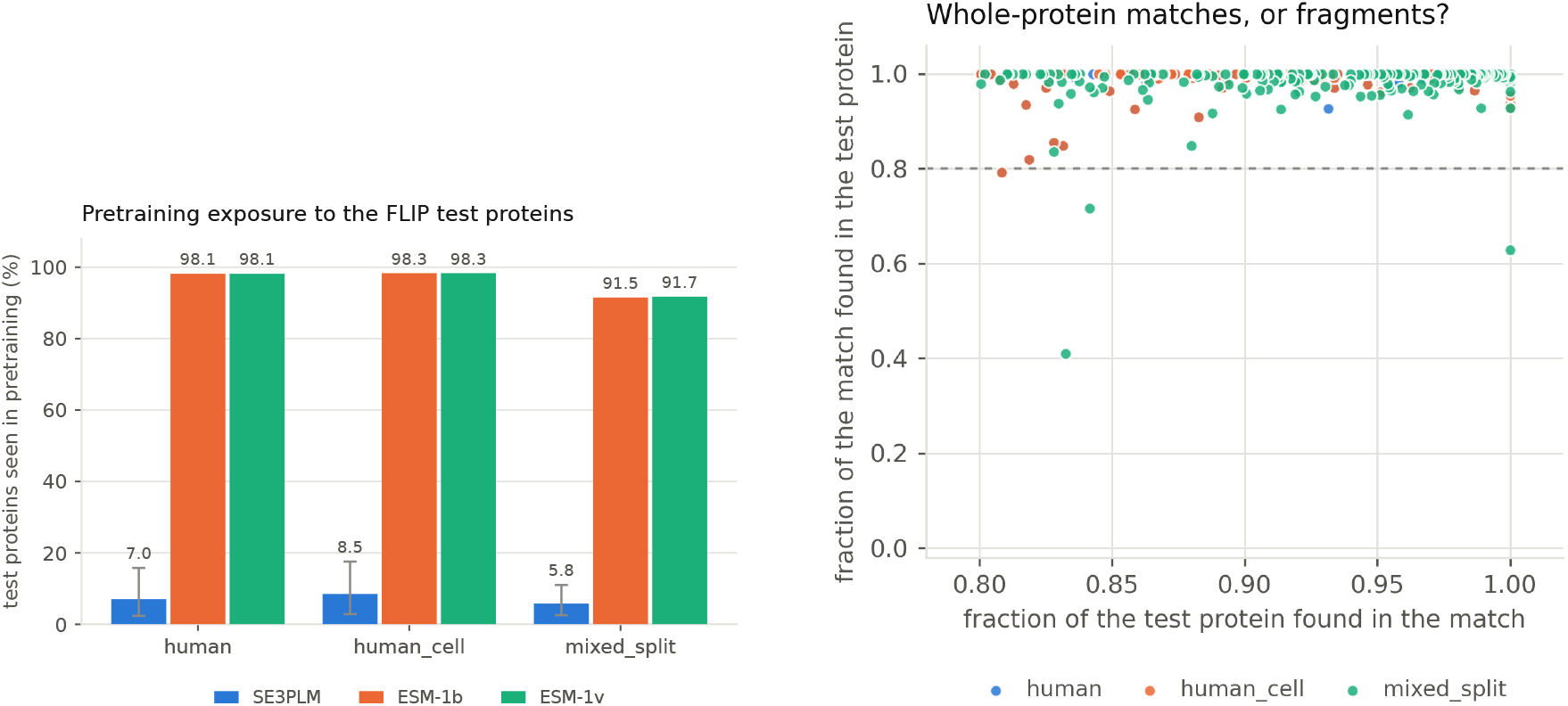
**Left:** Pretraining exposure with each model judged by its own criterion: BioT uses at least 0.95 of 3-mers in one pretraining protein; ESM uses UniProt membership at the clustering identity. Ours is the stricter criterion. **Right:** Each point is a test protein flagged at 0.80, placed by forward and reverse containment; points near (1, 1) are whole-protein matches.

**Figure 9.**
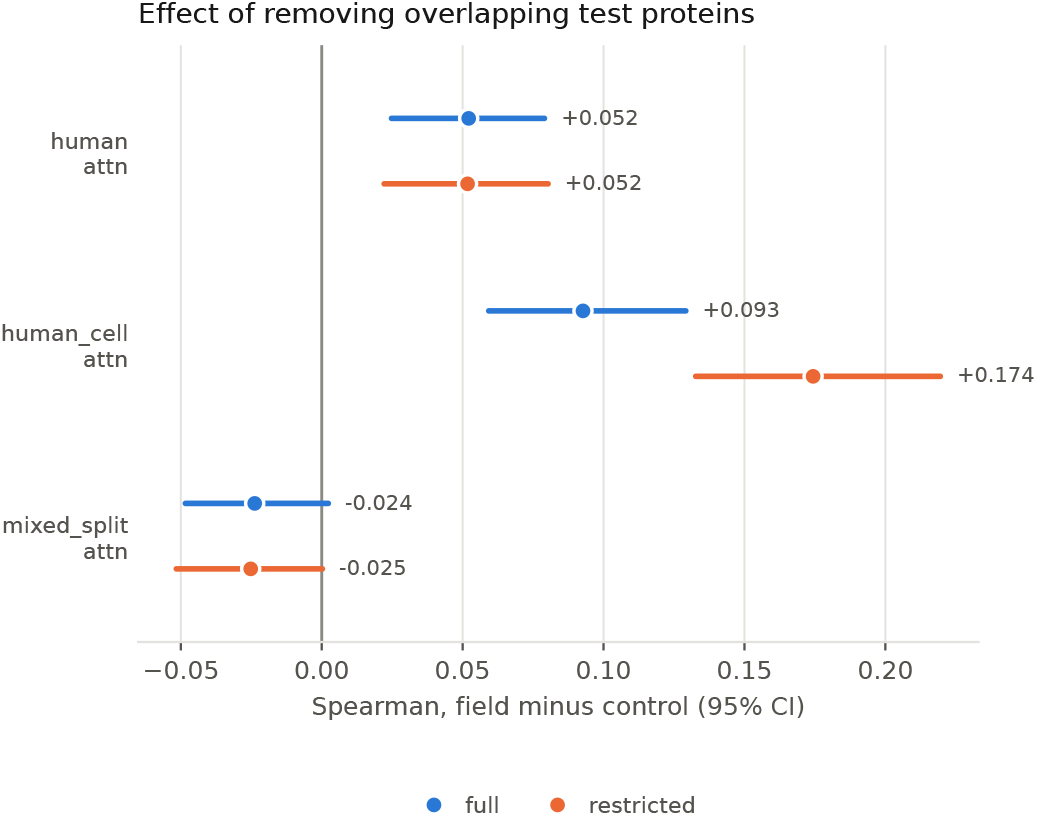
Composition-controlled difference, re-run. (attention head, point estimates with 95% intervals; values in Table 14).

**Figure 10.**
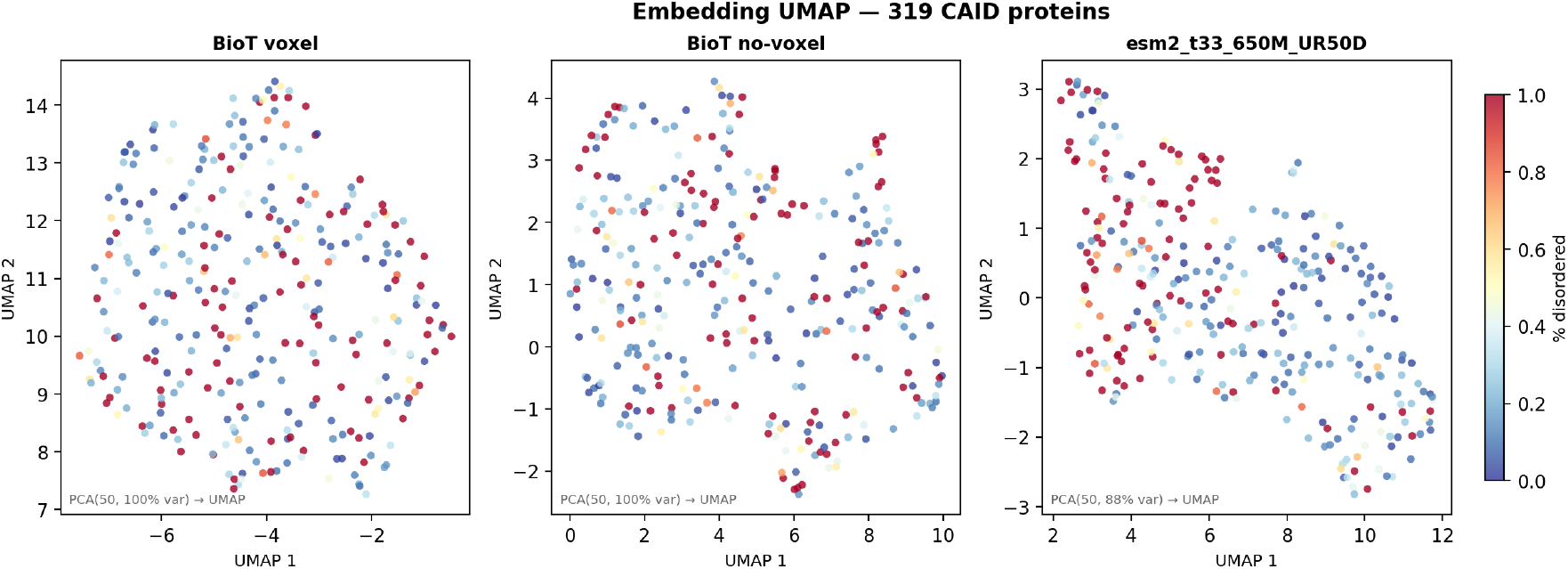
UMAP of mean-pooled per-protein embeddings on 319 CAID proteins, coloured by ground-truth disorder fraction (blue = ordered, red = disordered). PCA to 50 components precedes UMAP in all panels. BioT +field (*left*) shows tighter grouping by disorder fraction than the *−*field control (*centre*); ESM-2 (*right*) shows the clearest separation, consistent with its 91% pretraining overlap and 20*×* larger embedding dimension.

**Figure 11.**
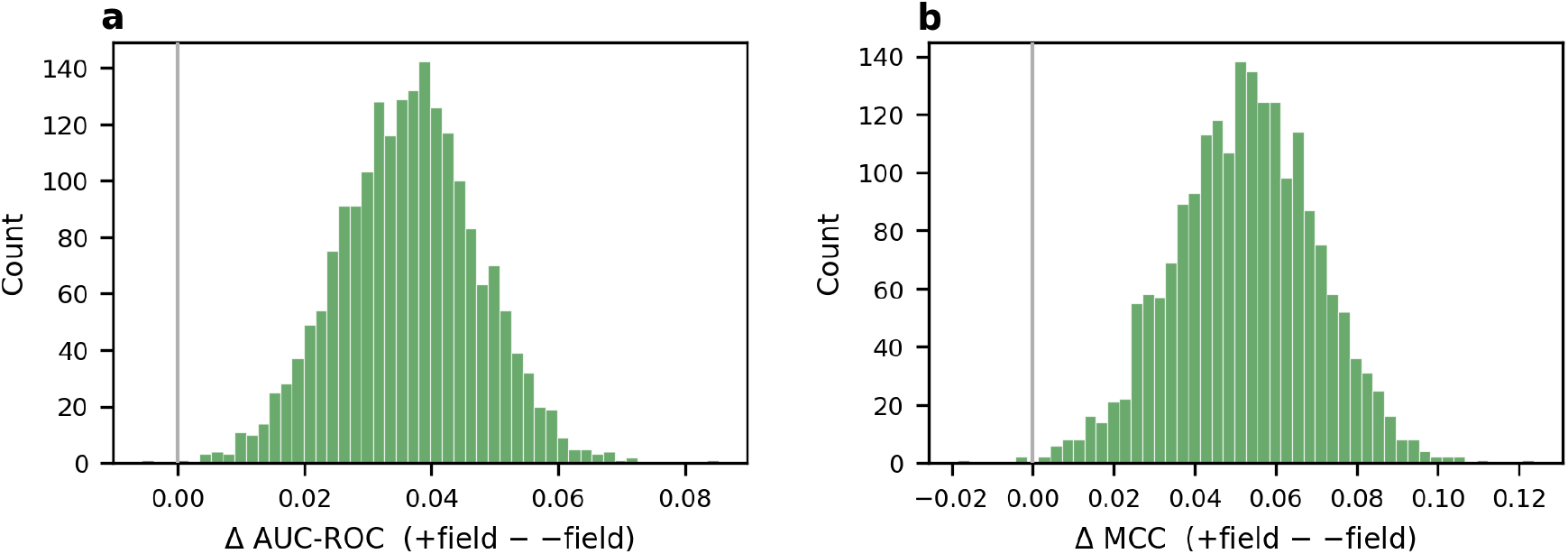
Paired bootstrap distributions for the CAID ablation. (2,000 replicates). *(a)* Δ AUC-ROC (+field minus *−*field); *(b)* Δ MCC. The grey vertical line marks zero. Both distributions lie entirely above zero, confirming that the voxel objective’s improvement is not an artefact of protein sampling.

Table 14 repeats the composition-controlled analysis with the flagged proteins removed. On `human`, the attention head moves from +0.0521 to +0.0518 and the CNN head from +0.0330 to +0.0294, both intervals still clear of zero. This is the result this appendix exists to establish. The `human_cell` attention head nearly doubles to +0.1744, far more than removing 8.5% of proteins should produce, so we treat that split’s attention result as unstable rather than as corroboration. Removal deletes 5.8% to 8.5% of each split, so the overlap-removed intervals are wider by construction. An interval that widens enough to touch zero has lost precision, not necessarily the effect. Finally, containment at the 3-mer level detects near-duplicates but not remote homologues sharing a fold and a thermostability profile while sitting well below 0.80 in sequence identity. That phylogenetic question is what Appendix M examines instead.

### L. Full Transfer Results

### M. Species Stratification

Of the 3,134 mixed-split test proteins, 2,887 (92.1%) match a UniProt entry for one of the 13 Meltome Atlas organisms; the remaining 247 are unlabeled and enter no group. Six organisms reach 200 proteins. No group shows a difference separable from zero in the field arm’s favor (Table 16). One is separable against it (*A. thaliana* at −0.108), but with six groups tested one clearing an uncorrected 0.05 happens 26% of the time under the null. The largest positive, *E. coli* at +0.103, does not exclude zero. The reason is power: an effect the size of `human`’s would reach at most 2.1 standard errors in the largest group.

**Table 15.** FLIP Meltome, Spearman. Δ is field minus sequence-only, computed before rounding. Trained heads are means *±*s.d. over 10 paired seeds; ridge probes and composition baselines are deterministic. The composition baselines do not depend on the encoder, so Δ is zero for that reason (—). The interval is a 95% paired bootstrap over test proteins; like the seed s.d. it takes no account of pretraining-run variance. Bold marks an interval excluding zero.

| Split | Probe | Sequence-only | + Field | Δ | 95% interval |
|---|---|---|---|---|---|
| Mixed<br>(*n*=3134) | AA composition | 0.3204 | 0.3204 | — | — |
|  | Dipeptide | 0.3920 | 0.3920 | — | — |
|  | Ridge (mean-pooled) | 0.3496 | 0.3718 | +**0.0222** | [+**0.0012**, +**0.0444**] |
|  | Ridge (BOS) | 0.2810 | 0.2966 | +0.0156 | [−0.0093, +0.0402] |
|  | Ridge (summary stat.) | 0.3717 | 0.3876 | +0.0159 | [−0.0059, +0.0386] |
|  | CNN head | 0.4371 ± 0.016 | 0.4369 ± 0.011 | −0.0002 | [−0.0192, +0.0193] |
|  | Attn head | 0.4564 ± 0.004 | 0.4628 ± 0.005 | +0.0064 | [−0.0118, +0.0253] |
| Human<br>(*n*=1945) | AA composition | 0.5392 | 0.5392 | — | — |
|  | Dipeptide | 0.5121 | 0.5121 | — | — |
|  | Ridge (mean-pooled) | 0.5655 | 0.5672 | +0.0017 | [−0.0130, +0.0156] |
|  | Ridge (BOS) | 0.4517 | 0.4686 | +0.0169 | [−0.0082, +0.0420] |
|  | Ridge (summary stat.) | 0.5502 | 0.5677 | +0.0175 | [−0.0033, +0.0384] |
|  | CNN head | 0.5734 ± 0.006 | 0.5977 ± 0.007 | +**0.0243** | [+**0.0112**, +**0.0373**] |
|  | Attn head | 0.5711 ± 0.006 | 0.5964 ± 0.005 | +**0.0253** | [+**0.0108**, +**0.0399**] |
| Human-cell<br>(*n*=1366) | AA composition | 0.5438 | 0.5438 | — | — |
|  | Dipeptide | 0.5099 | 0.5099 | — | — |
|  | Ridge (mean-pooled) | 0.5472 | 0.5560 | +0.0089 | [−0.0067, +0.0251] |
|  | Ridge (BOS) | 0.4416 | 0.4697 | +**0.0281** | [+**0.0013**, +**0.0542**] |
|  | Ridge (summary stat.) | 0.5287 | 0.5399 | +0.0112 | [−0.0118, +0.0348] |
|  | CNN head | 0.5463 ± 0.008 | 0.5689 ± 0.007 | +**0.0226** | [+**0.0086**, +**0.0379**] |
|  | Attn head | 0.5599 ± 0.004 | 0.5623 ± 0.007 | +0.0024 | [−0.0154, +0.0205] |

**Table 16.**
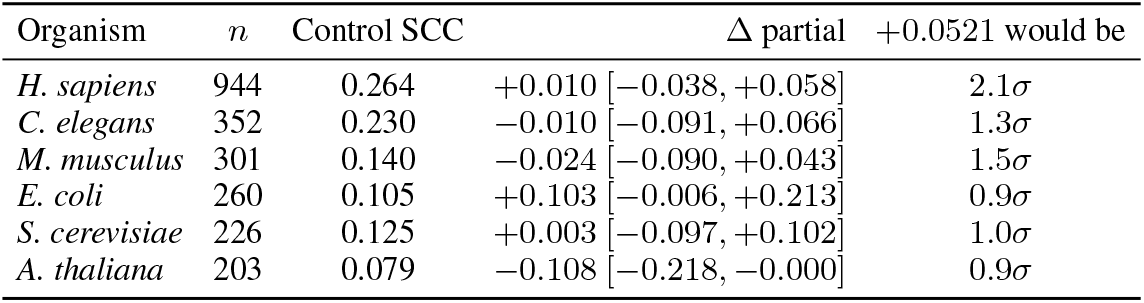
Field-minus-control difference in partial Spearman within single organisms of the mixed split, attention head, after residualizing the 400-dimensional dipeptide matrix. “Control SCC” is the sequence-only arm’s plain Spearman on that subset, for scale (it is 0.456 on the whole split). The last column is how many standard errors an effect the size of human’s +0.0521 would be in that group.

Within-organism thermostability is far less predictable than across organisms. The control’s Spearman falls from 0.456 on the whole split to between 0.079 and 0.264 within each of these six organisms, and the dipeptide baseline alone ranges from −0.140 to +0.178 against 0.392 on the whole split. Most of the mixed split’s predictability therefore lies between organisms rather than within them, which is the premise the phylogenetic explanation of Section 6 rests on.

### N. CAID Embedding Structure

Figure 10 shows UMAP projections of mean-pooled per-protein embeddings from BioT +field, BioT −field and ESM-2 (650M) on the 319 CAID disorder-PDB proteins, coloured by ground-truth disorder fraction. All three panels use PCA to 50 components followed by UMAP (BioT captures 100% of variance in 50 components; ESM-2 captures 88%, reflecting its higher intrinsic dimensionality).

ESM-2 organises the space most clearly: ordered proteins (blue) form one cluster and highly disordered proteins (red) another, with a smooth gradient between them. This is consistent with two advantages: 91% pretraining overlap (the model has seen nearly every test protein) and a 1280-dimensional embedding that gives the downstream probe ample signal.

BioT +field, despite its 64-dimensional embedding and 5% overlap, separates highly disordered proteins from ordered ones, with disordered proteins clustering toward the upper portion of the manifold. The −field control shows the same gross separation but with a noisier disorder gradient and more diffuse scattering of high-disorder proteins. The difference is qualitative, but it is consistent with the quantitative AUC-ROC and MCC improvements reported in Section 5.5: the voxel reconstruction objective organises the latent space in a way that correlates with structural disorder, even without explicit disorder supervision.

### O. CAID Bootstrap

Figure 11 shows the paired bootstrap distributions for the CAID ablation (2,000 replicates). In each replicate, the same proteins are resampled and scored by both BioT +field and BioT −field; the paired difference eliminates protein-level difficulty as a source of variance, yielding tighter intervals than marginal confidence intervals.

For Δ AUC-ROC (Figure 11a), the distribution is centred at +0.035 with a 95% interval of approximately [+0.007, +0.065]. For Δ MCC (Figure 11b), the distribution is centred at +0.054 with a 95% interval of approximately [+0.010, +0.102]. Neither distribution approaches zero, confirming that the voxel objective’s improvement on disorder prediction is not an artefact of protein sampling. The MCC distribution is wider than the AUC-ROC distribution, which is expected: MCC depends on a fixed probability threshold (0.5) and is therefore more sensitive to the calibration of the logistic regression probe.

## Notes

https://github.com/patjiang/Biophysical-Transformer

### Competing Interest Statement

The authors have declared no competing interest.

### Summary of Updates

Note on this revision. This version extends the original submission with three transfer experiments beyond FLIP Meltome: intrinsic disorder prediction on the CAID benchmark (Section 5.5), protein-protein interaction prediction on PLM-Interact (Section 5.6), and MHC I/II binding prediction (Section 5.7). Changes to the text include some added citations and revised discussion/future works notes. Code is now public at https://github.com/patjiang/Biophysical-Transformer.

## References

J. Abramson et al. Accurate structure prediction of biomolecular interactions with AlphaFold 3. Nature, 630:493–500, 2024. 10.1038/s41586-024-07487-w.

A. P. Bartók, R. Kondor, and G. Csányi. On representing chemical environments. Physical Review B, 87(18):184115, 2013. 10.1103/PhysRevB.87.184115.

M. Caron, H. Touvron, I. Misra, H. Jégou, J. Mairal, P. Bojanowski, and A. Joulin. Emerging properties in selfsupervised vision transformers. In IEEE/CVF International Conference on Computer Vision (ICCV), 2021. 10.1109/ICCV48922.2021.00951.

C. Dallago et al. FLIP: Benchmark tasks in fitness landscape inference for proteins. In NeurIPS 2021 Track on Datasets and Benchmarks (Round 2), 2021. Proceedings PDF. Preprint: 10.1101/2021.11.09.467890.

T. J. Dolinsky, J. E. Nielsen, J. A. McCammon, and N. A. Baker. PDB2PQR: an automated pipeline for the setup of Poisson– Boltzmann electrostatics calculations. Nucleic Acids Research, 32(Web Server issue):W665–W667, 2004. 10.1093/nar/gkh381.

A. Elnaggar et al. ProtTrans: Toward understanding the language of life through self-supervised learning. IEEE Transactions on Pattern Analysis and Machine Intelligence, 44(10):7112–7127, 2022. 10.1109/TPAMI.2021.3095381.

FLIP contributors. Stability dataset clustering data loss (issue 10). GitHub, J-SNACKKB/FLIP, 2022. https://github.com/J-SNACKKB/FLIP/issues/10.

M. Heinzinger et al. Bilingual language model for protein sequence and structure. NAR Genomics and Bioinformatics, 6(4):qae150, 2024. 10.1093/nargab/lqae150.

A. Jarzab et al. Meltome atlas—thermal proteome stability across the tree of life. Nature Methods, 17(5):495–503, 2020. 10.1038/s41592-020-0801-4.

J. Jumper et al. Highly accurate protein structure prediction with AlphaFold. Nature, 596:583–589, 2021. 10.1038/s41586-021-03819-2.

E. Jurrus et al. Improvements to the APBS biomolecular solvation software suite. Protein Science, 27(1):112–128, 2018. 10.1002/pro.3280.

S. Gelman, B. Johnson, C. R. Freschlin, A. Sharma, S. D’Costa, J. Peters, A. Gitter, and P. A. Romero. Biophysics-based protein language models for protein engineering. Nature Methods, 22(9):1868–1879, 2025. 10.1038/s41592-025-02776-2.

Z. Lin et al. Evolutionary-scale prediction of atomic-level protein structure with a language model. Science, 379(6637):1123–1130, 2023. 10.1126/science.ade2574.

Y. Liu et al. PLM-Interact: a protein language model for predicting protein–protein interactions. Nature Communications, 16:3520, 2025.

I. Loshchilov and F. Hutter. Decoupled weight decay regularization. In International Conference on Learning Representations (ICLR), 2019. https://openreview.net/forum?id=Bkg6RiCqY7.

J. Meier, R. Rao, R. Verkuil, J. Liu, T. Sercu, and A. Rives. Language models enable zero-shot prediction of the effects of mutations on protein function. In Advances in Neural Information Processing Systems, volume 34, pages 29287–29303, 2021. https://proceedings.neurips.cc/paper/2021/hash/f51338d736f95dd42427296047067694-Abstract.html.

S. K. Mylonas, A. Axenopoulos, and P. Daras. DeepSurf: A surface-based deep learning approach for the prediction of ligand binding sites on proteins. Bioinformatics, 37(12):1681–1690, 2021. 10.1093/bioinformatics/btab009.

S. Passaro et al. Boltz-2: Towards accurate and efficient binding affinity prediction. bioRxiv, 2025. 10.1101/2025.06.14.659707.

R. M. Rao, J. Liu, R. Verkuil, J. Meier, J. Canny, P. Abbeel, T. Sercu, and A. Rives. MSA Transformer. In Proceedings of the 38th International Conference on Machine Learning, volume 139, pages 8844–8856, 2021. https://proceedings.mlr.press/v139/rao21a.html.

A. Rives, J. Meier, T. Sercu, S. Goyal, Z. Lin, J. Liu, D. Guo, M. Ott, C. L. Zitnick, J. Ma, and R. Fergus. Biological structure and function emerge from scaling unsupervised learning to 250 million protein sequences. Proceedings of the National Academy of Sciences, 118(15):e2016239118, 2021. 10.1073/pnas.2016239118.

D. Sitkoff, K. A. Sharp, and B. Honig. Accurate calculation of hydration free energies using macroscopic solvent models. The Journal of Physical Chemistry, 98(7):1978–1988, 1994. 10.1021/j100058a043.

M. Steinegger and J. Söding. MMseqs2 enables sensitive protein sequence searching for the analysis of massive data sets. Nature Biotechnology, 35:1026–1028, 2017.

J. Su, M. Ahmed, Y. Lu, S. Pan, W. Bo, and Y. Liu. RoFormer: Enhanced transformer with rotary position embedding. Neurocomputing, 568:127063, 2024. 10.1016/j.neucom.2023.127063.

W. Torng and R. B. Altman. 3D deep convolutional neural net-works for amino acid environment similarity analysis. BMC Bioinformatics, 18:302, 2017. 10.1186/s12859-017-1702-0.

W. Torng and R. B. Altman. High precision protein functional site detection using 3D convolutional neural networks. Bioinformatics, 35(9):1503–1512, 2019. 10.1093/bioinformatics/bty813.

E. Wilson, J. K. Cava, D. Chowell, R. Raja, K. K. Mangalaparthi, A. Pandey, M. Curtis, K. S. Anderson, and A. Singharoy. The electrostatic landscape of MHC-peptide binding revealed using inception networks. Cell Systems, 15(4):362–373.e7, 2024. 10.1016/j.cels.2024.03.001.

J. Zhou, C. Wei, H. Wang, W. Shen, C. Xie, A. Yuille, and T. Kong. Image BERT pre-training with online tokenizer. In International Conference on Learning Representations (ICLR), 2022. https://openreview.net/forum?id=ydopy-e6Dg.

